# Pelagibacter, resolved

**DOI:** 10.64898/2026.04.03.716430

**Authors:** Torben N. Nielsen, Lauren M. Lui

**Author notes:** Both authors contributed equally.

## Abstract

*Pelagibacter*, the largest genus within the SAR11 clade, is the most abundant bacterium in the ocean, yet the vast majority of its species-level diversity remains uncharacterized at the genomic level. Here we present 135 complete *Pelagibacter* genomes — the largest such collection assembled to date — comprising 75 from Oxford Nanopore metagenomes of the San Francisco Estuary (SFE), 31 from a deeply sequenced station within the same transect, and 29 from public databases. These genomes define 52 species at 95% ANI, of which 44 (85%) are taxonomically novel. An expanded phylogeny incorporating 89 additional high-quality NCBI genomes confirms that our collection captures the phylogenetic backbone of the genus, with genomes from Hawaii, Namibia, and the Sargasso Sea nesting within SFE clades.

The pangenome is open (14,862 singletons, 62%), driven by two distinct mechanisms. First, a universal hypervariable region (HVR) at a conserved chromosomal position (7–15% from dnaA) is present in all 135 genomes, anchored by tRNA genes at both boundaries (Phe/His and Arg). The HVR carries genome-specific surface polysaccharide biosynthesis genes with a GC age gradient — highest GC at the tRNA boundaries, lowest in the center — consistent with a two-ended phage insertion model. Only this HVR is positionally conserved across the genus; the three other hypervariable regions previously described in a single reference genome are not. Second, scattered genomic islands throughout the chromosome contribute the remaining singleton content, including chimeric islands with genes from four bacterial phyla.

Biosynthetic pathway reconstruction reveals auxotrophies that are phylogenetically structured, not uniform: biotin, reduced sulfur, and glycine are genus-wide dependencies, while isoleucine, pantothenate, histidine, and glyoxylate cycle capacity vary across lineages with significant phylogenetic clustering. Structural annotation with ESMFold and Foldseek resolved 3,125 hypothetical proteins; 1,222 remain uncharacterized by any method, including a 47-amino-acid protein conserved in two-thirds of all genomes within a fixed operonic context — independently predicted by two gene callers yet matching nothing in any database. A controlled depth comparison at one station demonstrates that standard metagenome sequencing systematically underestimates *Pelagibacter* diversity, with three species recovered only at elevated depth and the species count at that station more than doubling (9 vs 4).

## Introduction

*Pelagibacter* is the largest genus within the SAR11 clade, the most abundant group of heterotrophic bacteria in the ocean, with a global population estimated at approximately 2.4 × 10²⁸ cells (Giovannoni, 2017). *Pelagibacter ubique* HTCC1062, the first cultivated member of the genus (Rappé et al., 2002), revealed the features that have come to define the lineage: unusually small genomes (∼1.3 Mbp), low GC content (∼29%), and coding densities exceeding 95%.

Despite their ecological dominance, the genomic diversity of *Pelagibacter* at the species level remains poorly characterized. The Genome Taxonomy Database (GTDB; Parks et al., 2022, 2025) currently lists hundreds of *Pelagibacter* species-level taxa, but the vast majority are represented only by fragmentary metagenome-assembled genomes (MAGs) or single-amplified genomes (SAGs). Complete, closed genomes — essential for accurate assessment of gene content, synteny, and mobile genetic elements — number only in the dozens across all public databases. This scarcity of complete reference genomes limits our ability to distinguish genuine biological variation from assembly artifacts, and to reconstruct complete metabolic pathways with confidence.

The difficulty of assembling complete *Pelagibacter* genomes from metagenomes is not merely a matter of sequencing depth. *Pelagibacter* species share substantial sequence similarity in core genes while co-occurring in the same environment, and each genome carries a hypervariable region of unique surface modification genes flanked by conserved sequence.

When multiple species are present, the conserved regions create ambiguities in assembly graphs while the species-specific HVR content creates branching paths that the assembler cannot resolve without sufficient coverage. Short-read metagenomics cannot resolve complete genomes under these conditions, and even long-read approaches require significant coverage to span the shared regions and resolve individual genomes.

Recent advances in long-read metagenomic assembly have begun to address this limitation. Current Oxford Nanopore Technologies (ONT) chemistry produces reads of sufficient length and accuracy for complete genome recovery directly from metagenomes, and assemblers such as myloasm (Shaw, Marin & Li, 2026) can exploit polymorphisms within long reads to resolve closely related genomes that would collapse with conventional approaches. The combination of deep ONT sequencing with assembly algorithms designed for metagenomic complexity creates an opportunity to recover complete *Pelagibacter* genomes at a scale that was previously impractical.

This study is the second in a series of companion papers characterizing the 20 most abundant bacterial genera in the SFE, each from complete genomes (biogeography paper in preparation). Here we exploit this opportunity in the San Francisco Estuary (SFE), a temperate, highly productive estuary where *Pelagibacter* populations span a salinity gradient from mesohaline to polyhaline waters (Lui & Nielsen, 2024; Lanclos et al., 2023; Rasmussen & Francis, 2023). From ONT metagenomes of 16 samples across 8 stations and 2 seasons, assembled with myloasm, we recovered 75 complete *Pelagibacter* genomes. An additional 31 complete genomes were recovered from one station where 4 additional flow cells were sequenced on top of the standard 2 (6 total, ∼3× depth), providing a controlled test of the relationship between sequencing depth and species recovery. Combined with 29 complete *Pelagibacter* genomes from NCBI (31 downloaded, 2 excluded after quality and taxonomic assessment), our collection of 135 genomes represents, to our knowledge, the largest set of complete *Pelagibacter* genomes ever assembled.

We use this collection to address three questions that require complete genomes and broad species-level sampling. First, how much *Pelagibacter* species diversity exists beyond what is currently described? Second, are the metabolic dependencies (auxotrophies) that characterize *Pelagibacter* universal features of the lineage, or do they vary systematically across clades — and if so, what does this imply about niche differentiation? Third, how much *Pelagibacter* diversity is missed by standard metagenomic sequencing, and what are the mechanistic reasons for this underestimation?

## Results

### Recovery of 135 complete *Pelagibacter* genomes

We collected 135 complete *Pelagibacter* genomes from three sources (Supplementary Table 1; Figure 1A). Seventy-five were assembled from ONT metagenomes of the SFE, sampled at 8 stations along the estuarine salinity gradient in summer (n = 8) and winter (n = 8). An additional 31 were assembled from a single deeply sequenced station (station 8, summer; hereafter “0S”), which combined the standard 2 flow cells with an additional 4, yielding approximately 3× the sequencing depth. Finally, 31 complete *Pelagibacter* genomes were downloaded from NCBI, encompassing all publicly available closed genomes for the genus at the time of analysis. Two were subsequently excluded: GCF_000195085 (*Ca.* Pelagibacter sp. IMCC9063), which belongs to genus IMCC9063 (SAR11 subclade II) and falls below skani’s (Shaw & Yu, 2023) ANI detection threshold (∼80%) against all other genomes in the dataset, and GCA_047724965, which showed an anomalously high fraction (18.5%) of truncated genes and was consistently absent from soft-core protein clusters.

**Figure 1A:**
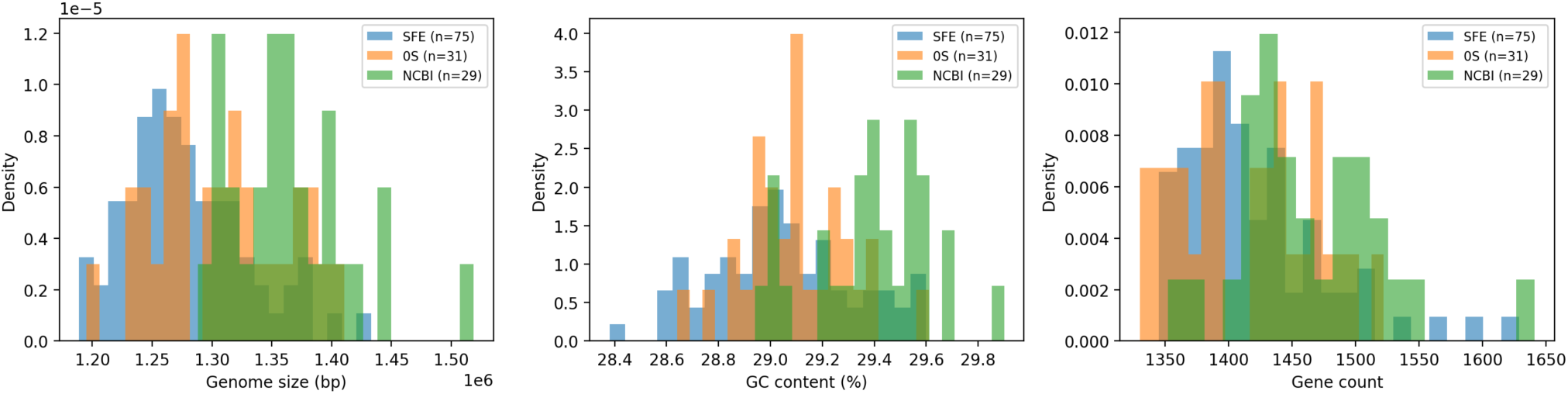
Distribution of genome sizes, GC content, and gene counts across the 135 genomes, colored by source.

All 135 genomes passed quality assessment with CheckM2 (Chklovski et al., 2023): all achieved ≥90% estimated completeness (mean 97.9 ± 1.7%) and <5% estimated contamination (mean 0.36 ± 0.55%). Genome sizes ranged from 1,188,876 to 1,518,778 bp (mean 1,304,203 ± 61,457 bp), consistent with the known range for *Pelagibacter*. GC content was uniformly low, ranging from 28.4% to 29.9% (mean 29.1 ± 0.3%). Gene calling with Pyrodigal in single-genome mode (Larralde, 2022) — important for the extreme AT-richness of *Pelagibacter* — identified 1,330 to 1,641 protein-coding genes per genome (mean 1,428 ± 59), with coding densities (fraction of genome length occupied by predicted CDS) of 95.9 ± 0.4%.

tRNA gene prediction with tRNAscan-SE v2.0.12 (Chan & Lowe, 2019) identified 30–36 tRNAs per genome (mean 32.1). All 135 genomes encode both the initiator fMet-tRNA and the elongator Met-tRNA (CAT anticodon). Cross-validation with ARAGORN v1.2.41 (Laslett & Canback, 2004) resolved one genome (M_u12081019) that tRNAscan-SE had scored as missing His-tRNA, bringing the count to 133 of 135 genomes with tRNAs for all 20 standard amino acids. The remaining two genomes each lack a single tRNA (7S_u40680507 missing His; 8S_u5554703 missing Lys), confirmed by both tools including tRNAscan-SE in maximum sensitivity mode (–max). These likely reflect wobble-pairing from a related anticodon rather than incomplete assembly. The uniformly complete tRNA complement provides independent evidence that these are genuinely complete genomes.

The SFE genomes were not uniformly distributed across samples. Summer stations contributed the majority (4S: 20 genomes; 7S: 18; 8S: 20), while winter stations contributed few (range 0–3).

### Fifty-two species, forty-four of which are novel

Pairwise ANI values were computed for all 135 genomes using skani (Shaw & Yu, 2023). Of the 9,045 pairwise comparisons, 8,860 yielded non-zero ANI values (range: 76.69–99.98%; mean among non-zero pairs: 82.15%). At the standard 95% ANI threshold for species delineation, the 135 genomes clustered into 52 species-level groups (Figure 1B; Figure 2A).

**Figure 1B:**
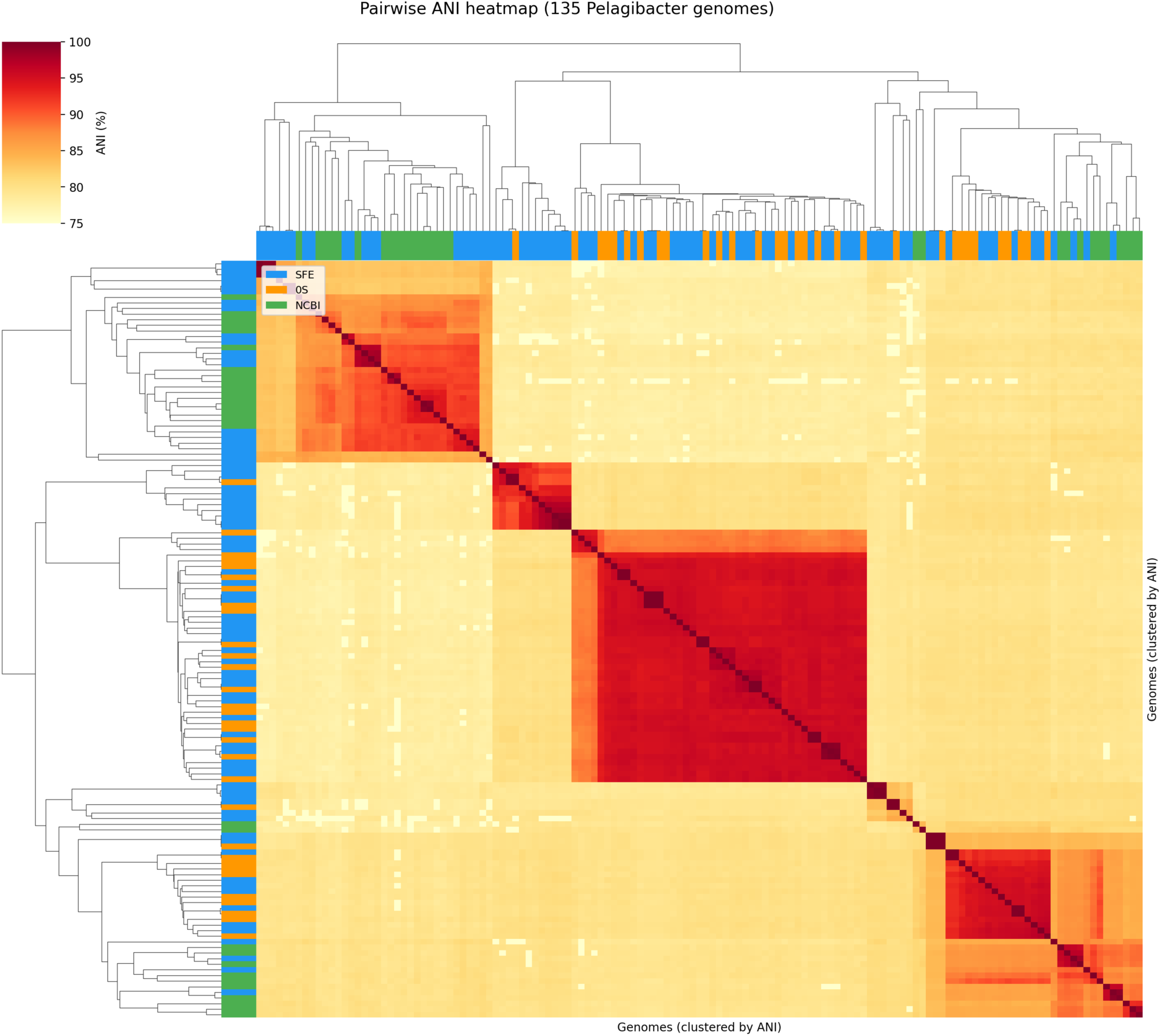
Pairwise ANI heatmap ordered by hierarchical clustering.

**Figure 2:**
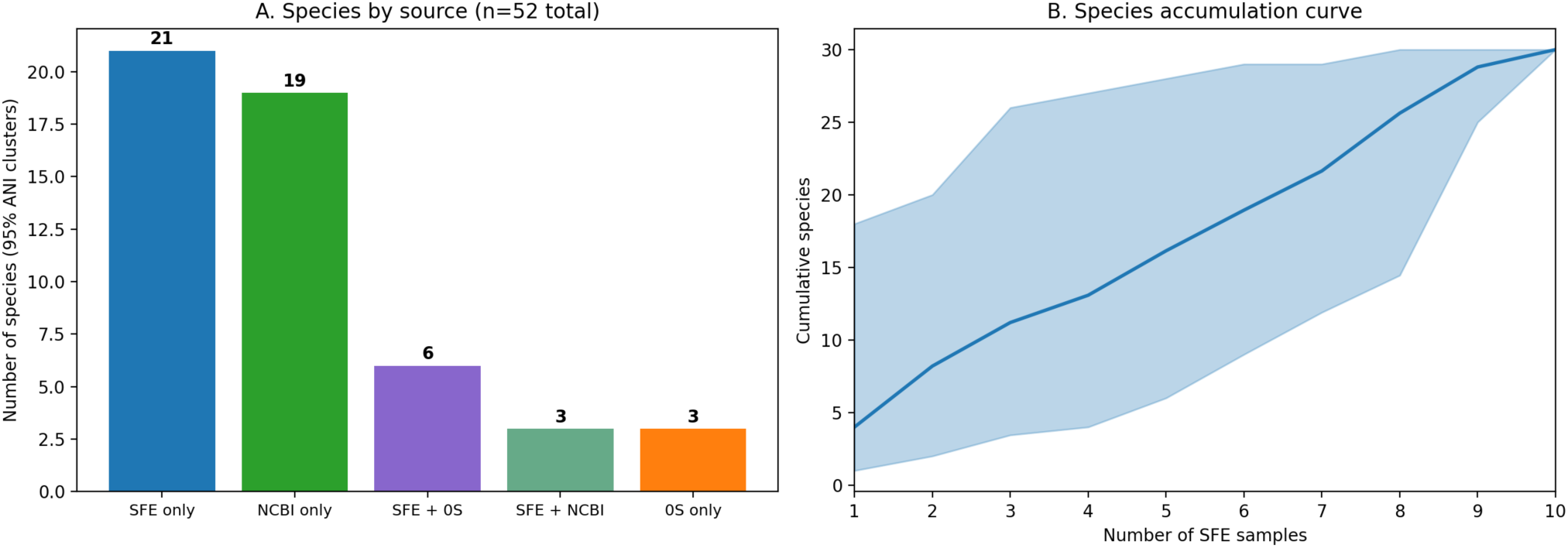
(A) Species cluster overlap among sources. (B) Species accumulation curve — permuted sample order, cumulative species count.

GTDB-Tk classification (Chaumeil et al., 2022; GTDB R226; Parks et al., 2025) assigned all 135 genomes to the genus *Pelagibacter*. Of the 52 species, 44 (85%) have no match to any named GTDB species and are therefore taxonomically novel. The remaining 8 correspond to established GTDB species: *Pelagibacter* sp008638165, *P.* sp028229795, *P.* sp001438335, *P.* sp902512765, *P.* sp943841085, *P.* sp007833635, *P. giovannonii*, and *P. ubique* (Supplementary Table 1).

Twenty-two of the 52 species included at least one NCBI genome, while 30 contained only SFE and/or 0S genomes. Among the 44 novel species, 26 have no prior genome in any public database; the remaining 18 include NCBI genomes that were previously deposited but remain taxonomically uncharacterized. In all cases, this study provides the first complete genomes for these species — the NCBI assemblies are predominantly fragmentary (scaffold or contig level). This novelty rate — 85% of species previously unnamed — underscores the magnitude of undiscovered *Pelagibacter* diversity, even in a single well-studied estuary (Figure 2B).

### Phylogenomic structure

Phylogenomic inference was performed on a supermatrix of 80 single-copy core protein clusters (present in exactly one copy in all 135 genomes; identified from the pangenome clustering described below), aligned with MAFFT and concatenated into 17,509 amino acid positions. IQ-TREE analysis with per-gene LG+G4+F partitions and 1000 ultrafast bootstrap replicates produced a well-resolved tree (log-likelihood −300,590.8; Figure 3): 68% of internal nodes (90/132) received ≥95% UFBoot support, 85% received ≥80%, and only 3 nodes (2%) fell below 50%. The tree was rooted using IMCC9063 (GCF_000195085) as an outgroup. IMCC9063 belongs to SAR11 subclade II, classified by GTDB as a separate genus (g IMCC9063) outside *Pelagibacter* despite being labeled “Candidatus Pelagibacter sp. IMCC9063” at NCBI. It was excluded from the pangenome analysis due to extreme divergence (below skani’s ANI detection threshold against all 135 genomes) but retains orthologs for 72 of the 80 core genes (mean 51% amino acid identity), providing a distant but alignable outgroup. The rooted tree was inferred from the same 80-gene supermatrix with IMCC9063 added to each alignment using MAFFT `--add ––keeplength`, yielding 136 taxa at 17,509 positions (13% gaps in the outgroup). The rooted tree (log-likelihood −323,997) shows comparable support to the unrooted tree: 67% of nodes ≥95% UFBoot, 85% ≥80%.

**Figure 3:**
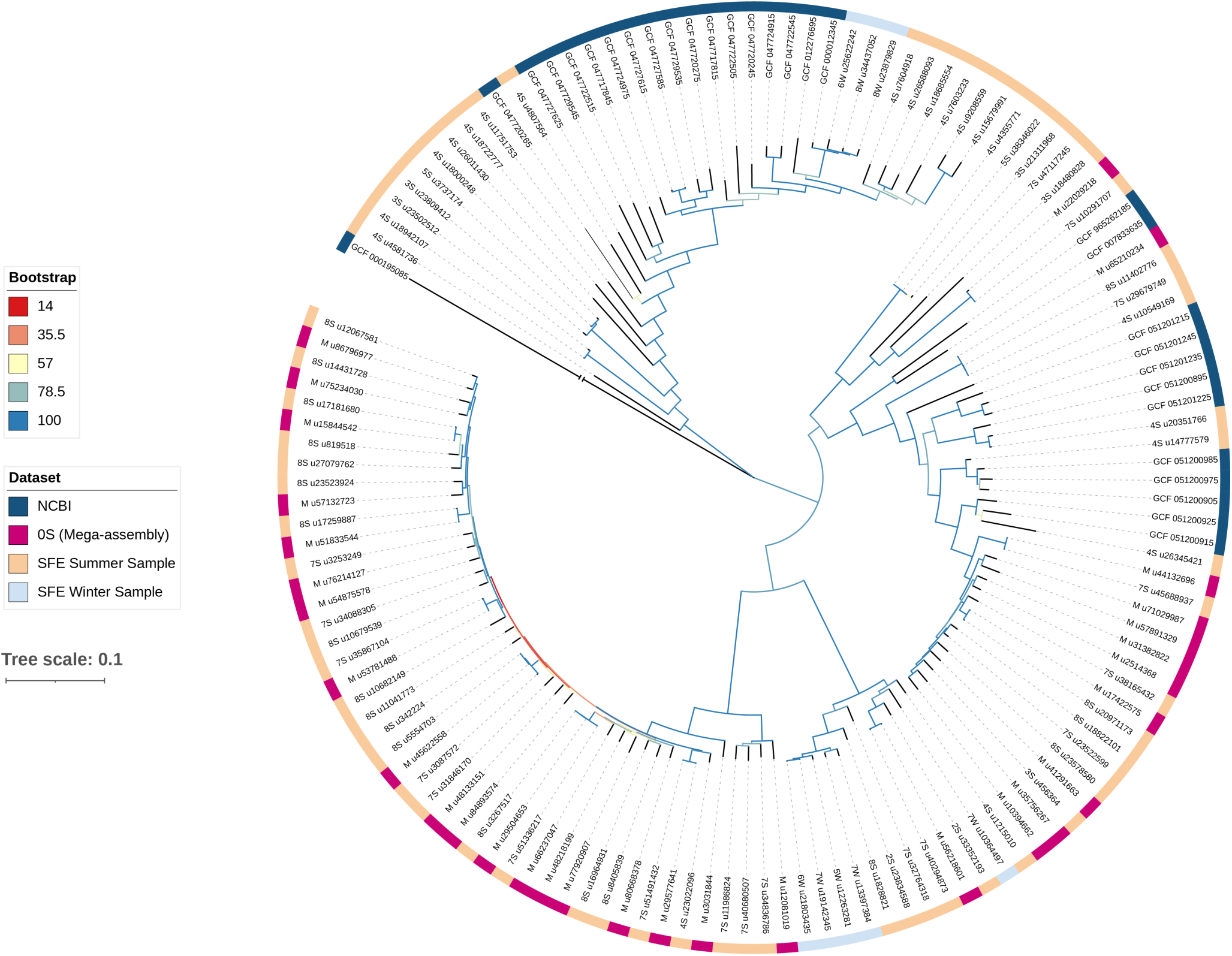
Rooted phylogenomic tree of 136 *Pelagibacter* genomes (135 ingroup + IMCC9063 outgroup). Branch colors indicate ultrafast bootstrap support (blue = 100, red = low). Outer ring indicates genome source: NCBI (dark blue), 0S mega-assembly (magenta), SFE summer samples (orange), SFE winter samples (light blue). Tree scale: 0.1 substitutions per site.

The tree resolves several deeply divergent lineages within *Pelagibacter*. Genomes from all three sources (SFE, 0S, NCBI) are interspersed throughout the tree rather than forming source-specific clades, indicating that the SFE harbors representatives of globally distributed *Pelagibacter* diversity, not just a locally divergent lineage. The 44 novel species are distributed across the tree, not confined to a single radiation. The NCBI genomes, which include isolates from diverse oceanic environments, are placed among SFE genomes at multiple points in the tree, confirming that the estuarine species are phylogenetically interleaved with oceanic diversity.

To test whether our collection captures the broader phylogenetic diversity of the genus, we constructed an expanded tree incorporating 89 additional high-quality *Pelagibacter* genomes from NCBI (≥90% completeness, <5% contamination, classified as g Pelagibacter by GTDB-Tk). Core gene orthologs were identified in each genome by searching against the 80 core cluster representatives, and alignments were extended using MAFFT `--add ––keeplength` to preserve the original alignment columns. IQ-TREE analysis of the resulting 225-taxon supermatrix (17,509 positions, 80 LG+F+G4 partitions, 1000 UFBoot replicates; log-likelihood −444,663) produced a tree with comparable support to the 135-genome tree: 70% of nodes ≥95% UFBoot, 83% ≥80%. The additional NCBI genomes — including isolates from Hawaii, Namibia, and the Sargasso Sea — nest within the clades defined by our SFE genomes rather than forming separate branches, confirming that our 135-genome collection captures the phylogenetic backbone of the genus (Figure S1).

### Pangenome architecture

MMseqs2 (Steinegger & Söding, 2017) clustering of 192,716 predicted proteins from all 135 genomes at 70% identity and 80% bidirectional coverage yielded 24,025 gene clusters (Figure 4A). Only 84 clusters (0.35%) were strictly core — present in all 135 genomes (Supplementary Table 2) — of which 80 occur in a single copy per genome (79 with full-length copies in all genomes plus one with a truncated copy in one genome; these 80 were used for phylogenomic inference, see above) and 4 carry paralogous copies in one or more genomes. An additional 406 clusters were soft-core (present in ≥128 of 135 genomes, ∼95%). The shell (2–94%) comprised 8,673 clusters, and 14,862 clusters (62%) were singletons found in only a single genome. Per-genome gene cluster counts ranged from 1,330 to 1,629 (Figure 4C), a 22% range despite all genomes sharing >80% ANI at the nucleotide level.

**Figure 4A:**
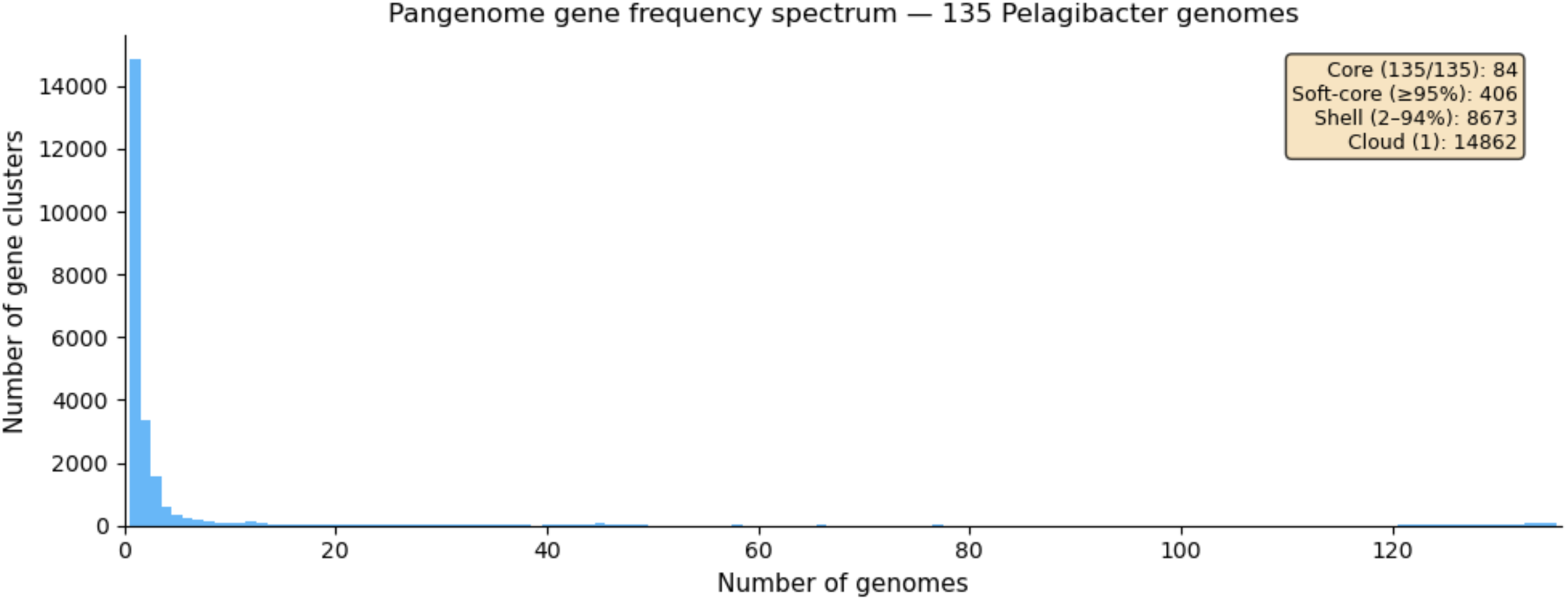
Gene frequency spectrum showing core/soft-core/shell/cloud distribution.

**Figure 4B:**
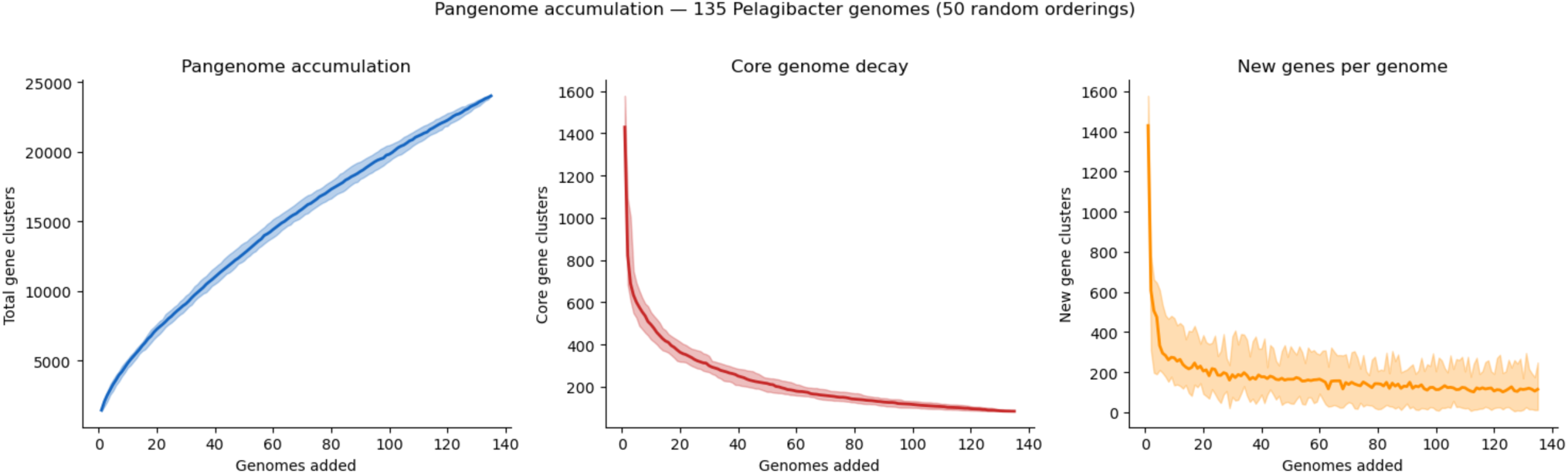
Pangenome accumulation curves — total clusters, core decay, and new genes per genome.

**Figure 4C:**
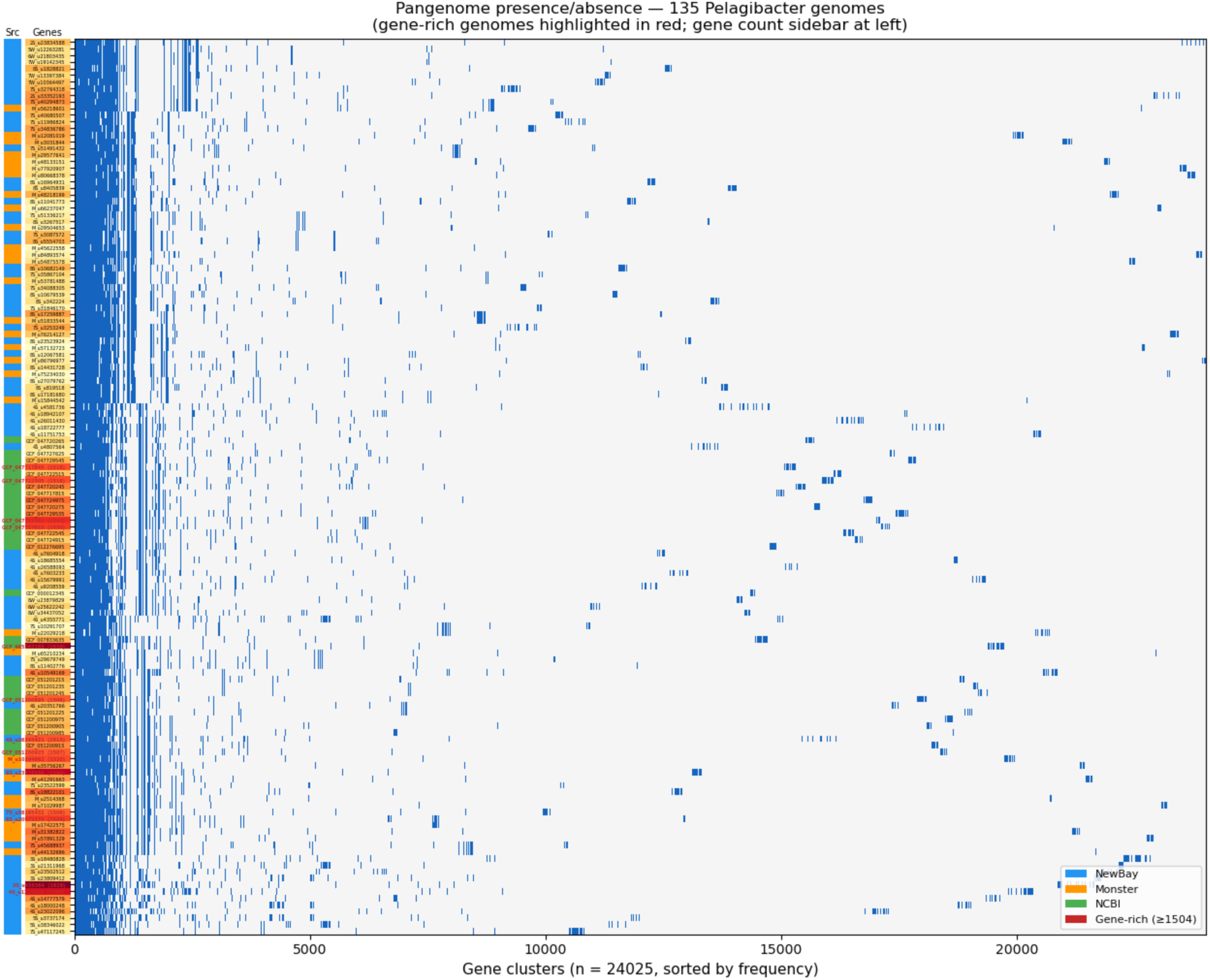
Phylogeny-ordered presence/absence heatmap with gene count sidebar.

**Figure 4D:**
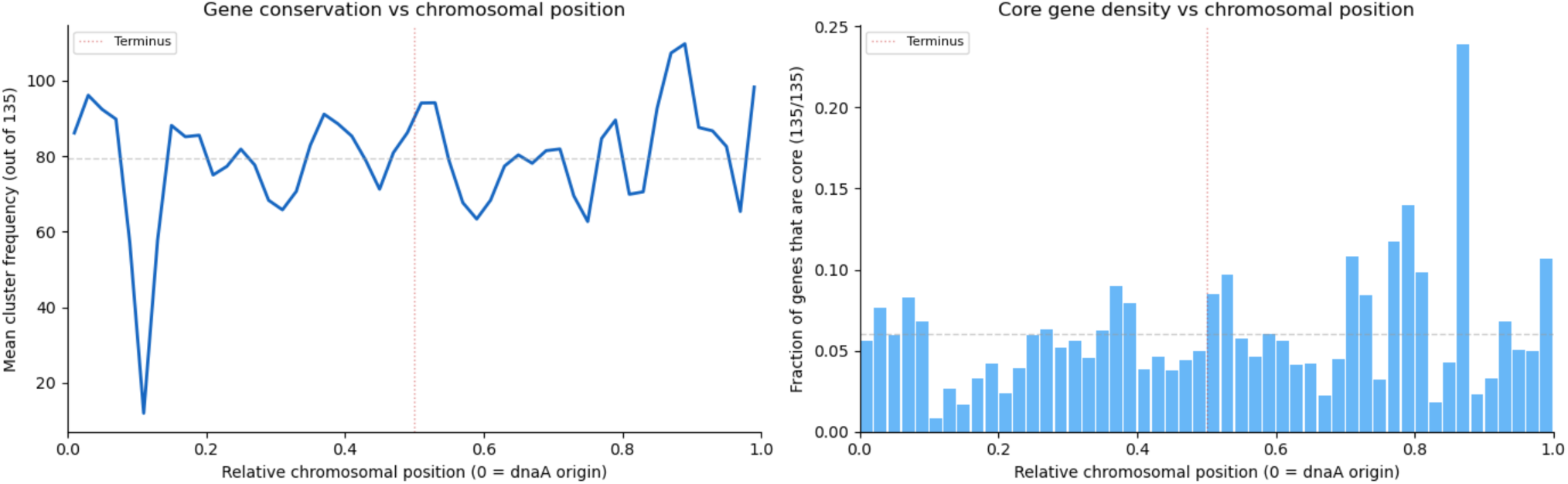
Gene conservation vs chromosomal position.

The pangenome is unambiguously open. Accumulation curves showed no plateau: each additional genome contributed a median of 115 new gene clusters even at 135 genomes (Figure 4B). The core genome decayed sharply in the first ∼30 genomes and stabilized near 84 clusters, indicating that the strict core is well-defined while the accessory genome continues to expand. This result is robust to the clustering threshold: at 50% identity the pangenome contains 17,318 clusters (57.9% singletons, ∼77 new per genome at 135), and at 90% identity it contains 45,076 clusters (60.9% singletons, ∼222 new per genome). The singleton fraction is stable across thresholds (58–62%), confirming that the open pangenome is not an artifact of the clustering parameters.

Assembly frameshifts are detectable and do not affect pangenome conclusions. ONT basecall errors in homopolymer runs can introduce indels that shift the reading frame, causing a single gene to be predicted as two adjacent ORFs or as one truncated ORF followed by an uncalled gap. We detected these frameshifts by two complementary methods: (1) adjacent short ORFs whose protein sequences both match the same UniRef90 target (split genes: 2,878 across 131 genomes), and (2) proteins significantly shorter than their UniRef90 match (query/ target length ratio <0.7) followed by an intergenic gap approximately matching the missing portion (truncated-with-gap: 669 across 123 genomes). In total, 3,547 genes (1.8%) are affected by frameshifts, concentrated in the same long essential genes — RNA polymerase beta’ (32 instances), FtsQ (22), DNA polymerase III alpha (16) — and in lower-depth assemblies (range 1–88 per genome, mean 26). Every frameshifted gene is still detected; the effect is overcounting (the 2,878 split genes inflate the ORF count by one each) rather than gene loss. No pathway step in the auxotrophy analysis is scored as absent due to frameshifts, and the pangenome size is inflated by at most 1.5%.

Gene content correlates with sequence divergence but does not explain the full range of variation. A Mantel test comparing pairwise gene content distance (Jaccard on the binary presence/absence matrix) against pairwise ANI distance yielded a Spearman correlation of ρ = 0.85 (p < 0.001; 9,999 permutations; ρ = 0.85 with or without the 185 pairs below skani’s detection threshold) — close relatives share more gene clusters than distant ones, as expected. However, this correlation is driven largely by the shared core and soft-core genome. Among the most gene-rich genomes, unique gene content (200–300 singletons per genome) varies independently of phylogeny: these genomes are scattered across the tree and do not share their unique gene complements with each other, consistent with independent acquisition of genomic islands rather than vertical inheritance.

A subset of genomes carry disproportionately large accessory gene inventories. The 14 most gene-rich genomes (≥1,506 clusters) each carry 200–300 unique gene clusters absent from all other genomes in the dataset. These gene-rich genomes are scattered across the phylogeny — they do not form a single clade — and their unique gene content is not shared with each other. This pattern is consistent with independent acquisition of genomic islands or phage-derived regions, rather than vertical inheritance of an ancestrally larger genome.

Pairwise comparison of accessory gene sharing among the gene-rich genomes revealed two types of structure: (1) closely related genome pairs that share nearly identical accessory content (e.g., GCF_047727585 and GCF_047727615 share 1,318 of ∼1,450 accessory clusters), reflecting recent common ancestry with the same mobile element complement; and (2) phylogenetically distant genomes that share 600–1,000 accessory clusters, suggesting a common pool of mobile or variable genes that recombines across species boundaries.

#### Singleton clusters are genomic islands, not annotation noise

The 14,862 singletons — clusters found in only one genome — are not spurious predictions or orphan fragments (Figure 5). The vast majority (92.6%) have UniRef90 hits, and their mean length (242 aa) is only modestly shorter than non-singletons (264 aa). Functionally, 54.5% are annotated housekeeping or metabolic genes with no close homolog in the other 134 genomes at the 70% identity clustering threshold. Cell surface modification genes (glycosyltransferases, O-antigen ligases, polysaccharide biosynthesis; 8.7%) and restriction-modification systems (5.8%) are enriched relative to the non-singleton pangenome, while phage structural genes and transposases are rare (0.3% combined). Singletons have lower GC content than non-singletons (25.3% vs 26.9%, a difference of 1.6 percentage points; Mann-Whitney U = 5.4 × 10⁷, p < 10⁻¹⁶⁰; n = 14,862 vs 9,163), and both are below the genome-wide average of 29.1%, which includes rRNA, tRNA, and intergenic sequences that elevate the overall GC relative to protein-coding regions; 51% of singletons deviate from the genome mean by more than 5 percentage points, a signature of recent horizontal acquisition.

**Figure 5:**
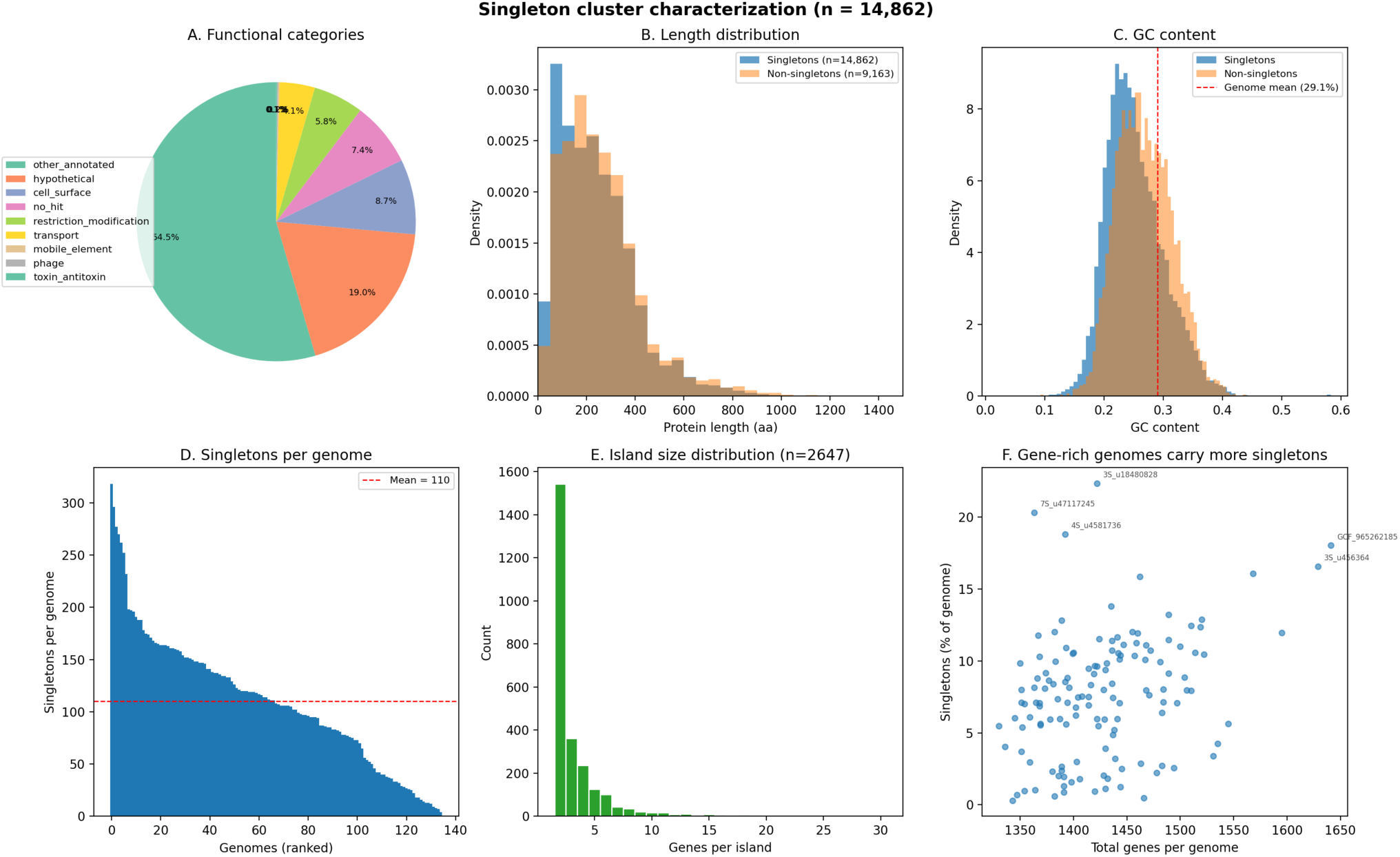
Singleton cluster characterization. (A) Functional categories. (B) Length distribution vs non-singletons. (C) GC content — singletons shifted below genome mean. (D) Singletons per genome. (E) Genomic island size distribution. (F) Gene-rich genomes carry more singletons.

Most singletons are not scattered individually but cluster in genomic islands: 77.7% occur in runs of ≥2 adjacent singletons, with 2,647 islands detected across the 135 genomes (mean 4.4 genes, up to 100 genes). These islands are the physical units of gene content variation — discrete insertions carrying surface modification, defense, and metabolic cargo on a background of conserved chromosomal architecture. Per-genome singleton counts range from 4 to 318 (mean 110 ± 64, or 7.7% of genes). The most gene-rich genomes are gene-rich specifically because they carry more and larger islands, not because they have duplicated core functions.

This pattern — islands of low-GC, HGT-acquired genes carrying surface and defense functions, inserted into a conserved chromosomal backbone — is consistent with the prophage-mediated gene transfer described in SAR11 by Morris et al. (2020), who showed that SAR11 prophages — members of the abundant pelagiphage group first described by Zhao et al. (2013) — spontaneously induce at low rates, maintaining lysogeny while enabling horizontal transfer of cargo genes. The near-absence of phage structural genes among our singletons (0.1%) suggests that the structural components are lost or degraded after integration, while the ecologically useful cargo — surface modification, restriction-modification, and metabolic genes — persists.

#### Taxonomic origins of horizontally acquired genes

The UniRef90 hit descriptions include the source taxonomy, enabling us to trace the phylogenetic origin of singleton genes. The majority (81%; 11,115/13,755) have best hits to other *Pelagibacter* or SAR11 sequences — genes present elsewhere in the genus but too divergent to cluster at the 70% identity threshold used in our pangenome analysis. Among the remainder, 38 singletons have best hits to non-*Pelagibacter* organisms at ≥70% amino acid identity, indicating recent cross-lineage acquisition. (Best-hit taxonomy identifies the closest sequenced relative of the donor, not necessarily the direct donor; the true transfer route may involve intermediate hosts or phage vectors.)

Individual islands reveal mosaic origins. A six-gene polysaccharide biosynthesis island in GCF_047729545 contains genes whose closest UniRef90 matches span four bacterial phyla: an NAD-dependent epimerase matching *Parabacteroides distasonis* (Bacteroidota; 80.9% identity), an NAD-dependent epimerase matching *Leptospira santarosai* (Spirochaetota; 63.9%), a nucleotide sugar dehydrogenase matching *Lacinutrix* sp. (Bacteroidota; 77.5%), a Vi polysaccharide biosynthesis gene matching *Pelagibacter* (Alphaproteobacteria; 69.0%), a nucleotide sugar dehydrogenase matching *Prochlorococcus marinus* (Cyanobacteria; 40.3%), and a sulfotransferase matching *Pelagibacter* (100%). This chimeric island is functionally coherent (all genes involved in surface polysaccharide biosynthesis) but phylogenetically mosaic, with closest relatives spanning four phyla — whether these genes were acquired individually or arrived as a pre-assembled unit cannot be determined from the data.

A DUF4910 domain-containing gene family illustrates a different mode of exchange. This gene is present in 19 of 135 genomes at widely varying sequence identities: four NCBI genomes carry copies at 100% identity to other *Pelagibacter* references, five genomes carry copies whose closest matches are to *Prochlorococcus* (79–84% identity), and the remainder match divergent *Pelagibacter* sequences (45–67%) or other marine bacteria including *Desulfovibrio* and *Aeromonas*. In one genome (8S_u10682149), the gene is split into two adjacent ORFs by a frameshift — likely an assembly-induced homopolymer indel — while in another (8S_u342224) it is intact at 430 aa and 83.6% identity to the *Prochlorococcus* version. The presence of this gene family at such disparate identities across both *Pelagibacter* and *Prochlorococcus* — the two most abundant bacterial lineages in the ocean — suggests ongoing gene exchange between co-occurring populations, rather than a single ancestral transfer event.

#### A universal hypervariable region dominates the accessory genome

The largest singleton islands — up to 100 consecutive singleton genes spanning 59 kbp — are not randomly positioned. The ten largest islands across all genomes occupy the same chromosomal locus: 7–15% of the genome from dnaA (∼115–200 kbp). This corresponds to the hypervariable region HVR2 identified by Wilhelm et al. (2007) in metagenomic fragments from the Sargasso Sea. Wilhelm et al. described four hypervariable regions in HTCC1062, the largest being HVR2 at 48 kb. With 135 complete genomes, we can now assess which of these are genus-wide features: only HVR2 is positionally conserved. No other chromosomal window shows comparable singleton enrichment (3,843 singleton genes in the 10–15% window vs 741 in the next highest). The other three HVRs described in HTCC1062 do not correspond to genus-wide hotspots, and we found no evidence that they are conserved features beyond that single genome.

Every genome carries genes at the HVR locus (99–116 genes in the 7–15% window), but the gene content is almost entirely genome-specific: 124 of 135 genomes have singleton islands here (median 36 singleton genes, range 2–102). The 11 genomes without singleton islands in the HVR still carry genes at this position, but their variants happen to cluster with close relatives at the 70% identity threshold rather than being unique.

The HVR gene content is dominated by surface polysaccharide biosynthesis: glycosyltransferases, NAD-dependent epimerases, DegT/DnrJ/EryC1/StrS family aminotransferases, GDP-mannose dehydratases, and sulfotransferases. These are the enzymes that build and modify the cell surface carbohydrate structures that phages use as receptors. The GC content of HVR singleton genes is markedly depressed (mean 25.3% vs 29.1% genome average), consistent with acquisition from diverse external sources. Many genes have best UniRef90 hits to marine metagenome sequences or to distantly related organisms across multiple phyla, indicating that the HVR accumulates horizontally acquired surface modification genes from a broad donor pool.

Wilhelm et al. (2007) reported that the HVR in HTCC1062 is flanked by rRNA genes. With 135 complete genomes, we can now assess the full spatial structure of this region. Infernal cmsearch against bacterial 16S, 23S, and 5S covariance models reveals that every genome has a single copy of each rRNA gene (one genome has two 23S copies). The 16S and 23S are adjacent (median spacer 477 bp, the ITS region), but the 5S is separated from the 16S–23S pair by a median of 46,805 bp — the rRNA genes do not form a single operon. In 130 of 135 genomes, the 16S–23S pair sits at 8–10% from dnaA and the 5S at 12–14%, both within the HVR. The remaining 5 genomes — belonging to two sister species (sp0191, 3 genomes; sp0298, 2 genomes) from three independent samples — have the 16S–23S at ∼79% and the 5S at ∼75%, on the opposite replichore. All 5 are low-depth (depth1 8–13×) but circular and HQ, and the consistency of the rearrangement across two species points to a single ancestral translocation rather than an assembly artifact.

The HVR is thus bounded by tRNA genes, with the rRNA pair as an internal landmark. A tRNA cluster is conserved at the left (origin-proximal) HVR boundary: Phe-tRNA and His-tRNA are present at the 5–7% position in 130 and 132 of 135 genomes, respectively. A second tRNA cluster (Arg-tRNA) is conserved at the right (terminus-proximal) boundary in 79 of 135 genomes at 14–17%. The 16S–23S pair sits between the left and right tRNA boundaries, and the 5S sits near the right boundary. The full spatial order in 130/135 genomes is: dnaA (0%) → Phe/His-tRNA (5–7%) → HVR singleton content → 16S–23S (8–10%) → more HVR content → 5S (12–14%) → Arg-tRNA (14–17%). Wilhelm et al.’s observation that rRNA genes flank the HVR was correct for HTCC1062, but with 135 genomes it is clear that the tRNA genes are the conserved structural anchors while the rRNA genes — which are not a complete operon — occupy a conserved but internal position. The 5 genomes where the rRNA pair has relocated retain the tRNA boundaries at the same positions, confirming that the tRNAs, not the rRNAs, define the HVR.

tRNA genes are classic phage integration sites in bacteria, and their conservation at the HVR boundary across a genus with otherwise extensively rearranged gene order suggests a mechanistic explanation for the HVR’s positional stability: phages integrate at the conserved tRNA locus, deposit surface modification cargo genes, and the phage structural genes subsequently degrade — leaving the cargo behind as the HVR content. Why these tRNAs remain positionally fixed while most other operonic blocks shuffle freely is unclear; their proximity to dnaA (∼5–7% of the chromosome from the origin) may place them in a region with too few intervening operon boundaries to permit rearrangement. This model is consistent with the prophage dynamics described by Morris et al. (2020), where SAR11 prophages spontaneously induce at low rates, and with the near-absence of phage structural genes among HVR singletons despite their clear HGT signatures.

The internal structure of the HVR supports a two-ended insertion model. Across 4,684 HVR singleton genes in 132 genomes, GC content decreases from 25.5% near the tRNA boundaries to 22.3% in the center of the HVR (Figure 6; Spearman ρ = −0.17 for GC vs position, p < 10⁻³²), and GC deviation from the genome mean correspondingly increases toward the center (ρ = +0.16, p < 10⁻²⁹). UniRef90 sequence identity shows a U-shaped profile: highest at both boundaries (72% and 76%) and lowest in the center (61–63%). Both patterns are consistent with a model in which new genes are inserted at the tRNA-anchored boundaries and the oldest, most diverged cargo accumulates in the center — furthest from either integration point, with the most time to diverge from its donor and the most exposure to the host’s AT-biased mutational pressure. Analogous age gradients have been described in integron cassette arrays, where cassettes nearest the integrase are newest (Mazel, 2006), but a two-ended gradient within a chromosomal hypervariable region has not, to our knowledge, been previously reported.

**Figure 6:**
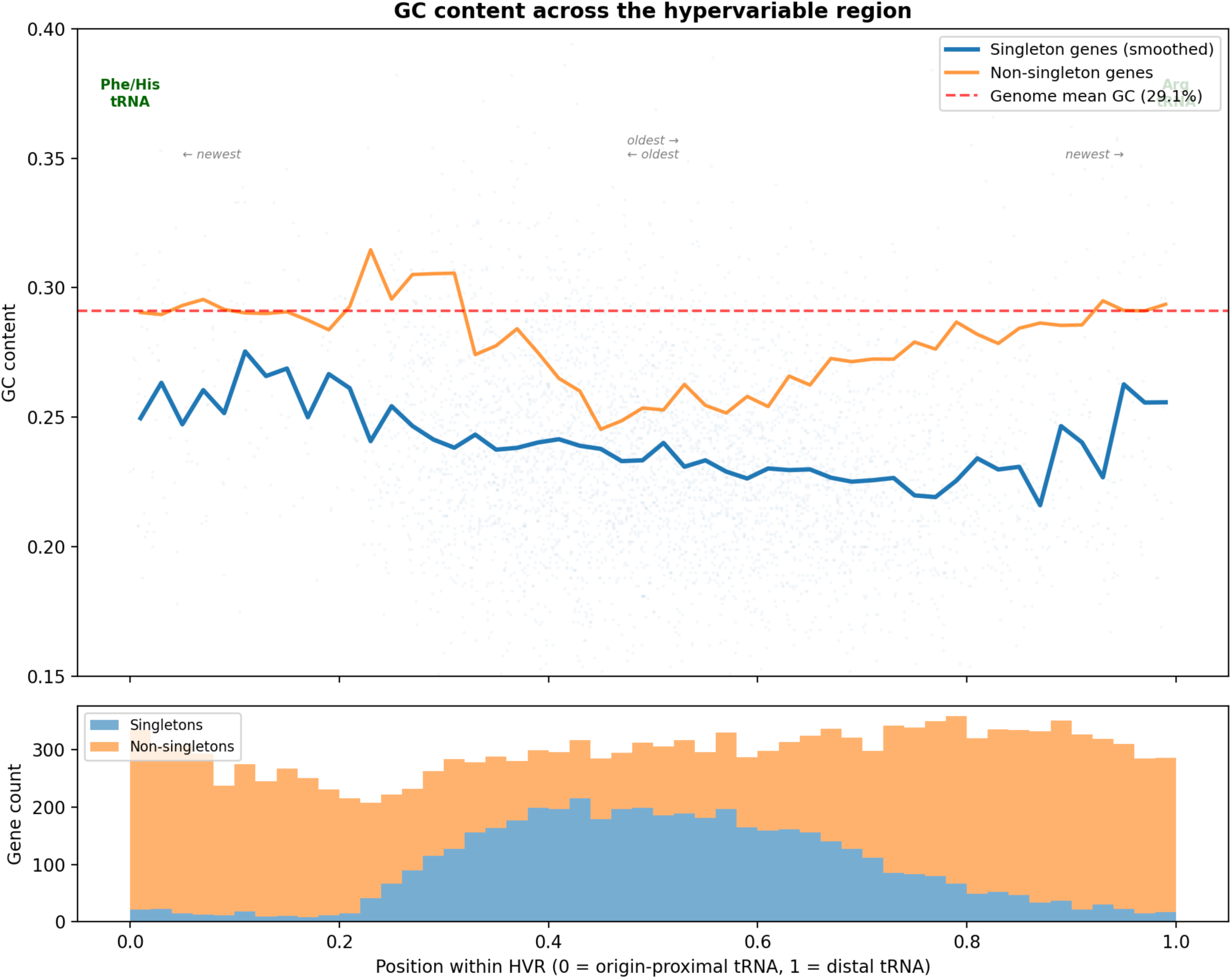
GC content across the hypervariable region. Top: singleton genes (blue) and non-singleton genes (orange) show a bathtub curve — highest GC at the tRNA-anchored boundaries, lowest in the center. Red dashed line: genome mean (29.1%). Bottom: number of genes at each position across 132 genomes, colored by singleton status. See also Figure S2 for GC deviation and UniRef90 identity scatter plots.

The HVR model is consistent with the prophage dynamics described for SAR11. Our filtration protocol (0.1 µm filters allowed to clog; Lui & Nielsen, 2024) captures both bacterial cells and free phage particles in the same DNA extraction, and geNomad (Camargo et al., 2024) identified 32,000–128,000 viral contigs per sample. Searching HVR singleton proteins against viral proteins from the same sample confirmed that most HVR genes have homologs in the co-occurring viral fraction; however, a control search of non-HVR conserved genes against the same viral proteins yielded comparable match rates (62% vs 80% at e-value 10⁻⁵), indicating that the match rate reflects general host–phage sequence sharing rather than HVR-specific enrichment. The extensive prophage content in our samples — consistent with the spontaneous induction dynamics described by Morris et al. (2020) — means that phage-carried copies of housekeeping and surface modification genes coexist with their chromosomal counterparts in the same metagenome. The tRNA-anchored boundaries, depressed GC, and phylogenetically mosaic gene content of the HVR remain strong indirect evidence for phage-mediated insertion, even though direct enrichment of viral matches in the HVR was not observed.

This finding resolves a tension in the pangenome data. The 14,862 singleton clusters and the open pangenome could, in principle, reflect a genome where variable genes are scattered uniformly. Instead, a large fraction of the singleton content is concentrated in a single positional hotspot consistent with a region maintained by phage-driven diversifying selection. The HVR is likely the physical locus where *Pelagibacter* generates surface antigen diversity, consistent with a model in which diversifying selection driven by pelagiphages (Zhao et al., 2013) maintains variant surface structures across co-occurring genomes.

However, the HVR does not fully account for the open pangenome. Of the 14,862 singletons, only 4,684 (31.5%) fall within the HVR; the remaining 10,178 (68.5%) are distributed across the rest of the chromosome. Recalculating the accumulation curve after excluding all HVR singletons, each additional genome still contributes ∼78 new clusters at 135 genomes (vs ∼115 with the HVR included), with little sign of saturation. The *Pelagibacter* pangenome is thus driven by two distinct mechanisms: phage-mediated surface coat diversification concentrated in the HVR, and independent genomic island insertions carrying metabolic, defense, and transport genes throughout the rest of the chromosome.

Gene conservation shows no positional bias along the chromosome (Figure 4D). Because all genomes were reoriented to begin at dnaA, we could test whether origin-proximal genes are more conserved than terminus-proximal genes — a pattern observed in larger bacterial genomes where multifork replication creates a gene dosage gradient. In *Pelagibacter*, genes near the origin were present in a mean of 84.3 of 135 genomes — virtually identical to the mean of 84.3 near the terminus — and the fraction of core genes showed only a marginal difference (6.9% vs 6.2%). The ∼1.3 Mbp *Pelagibacter* genome is evidently too small for replication-linked gene dosage effects to shape gene conservation patterns; instead, the spatial distribution of conserved and variable genes appears stochastic, consistent with mobile element insertion and excision as the dominant force shaping accessory gene content. The one exception is a visible dip in mean gene conservation at 8–10% from dnaA (Figure 4D), corresponding precisely to the HVR and its embedded 16S–23S rRNA genes — an independent confirmation of the HVR’s chromosomal position.

### Functional annotation and metabolic capacity

All 192,716 predicted proteins were searched against UniRef90 (UniProt Consortium, 2024) using MMseqs2 (e-value ≤ 10⁻⁵, top hit per query). Of these, 188,355 (97.7%) returned a hit, with 70.3% at ≥90% amino acid identity and 92.2% at ≥70% identity — indicating that the vast majority of *Pelagibacter* proteins are closely related to characterized sequences. Only 4,361 proteins (2.3%) had no detectable homolog in UniRef90. Among annotated proteins, 90.6% matched a functionally characterized UniRef90 cluster, while 9.4% were annotated as hypothetical or contained only domains of unknown function.

Length comparisons between query and target sequences provide an independent check on assembly quality. Of 188,355 annotated proteins, 92.6% had query/target length ratios between 0.8 and 1.2, consistent with full-length predictions. Only 6.8% (12,938) were shorter than 80% of their UniRef90 match, a fraction that includes both divergent members of broad protein families and potential assembly-induced frameshifts in AT-rich homopolymer regions. Mean query coverage was 0.99 and mean target coverage 0.96, indicating near-complete alignments.

The most frequently recovered UniRef90 entries correspond to universally conserved housekeeping genes: enoyl-ACP reductase (fatty acid synthesis; 142 hits across 135 genomes, reflecting paralogy), RNA polymerase alpha subunit (141), and ATP synthase beta subunit (140). These are consistent with the core genome composition and provide an independent validation of the protein clustering results.

KEGG ortholog assignments were made using KofamScan (Aramaki et al., 2020) with all KEGG profiles. Of 192,716 proteins, 111,630 (57.9%) received at least one significant KO assignment at the default threshold, spanning 1,171 unique KOs. However, KofamScan’s adaptive score cutoffs — calibrated across broad bacterial diversity — are too stringent for *Pelagibacter*’s divergent proteins. Among the 37,102 proteins that scored below the strict threshold but above 0.50×, the score/threshold ratio distribution is smooth with no natural break. We adopted a relaxed threshold of 0.75×, which increased the annotated fraction to 132,427 proteins (68.7%). The choice of 0.75× is not critical: repeating the pathway analysis at 0.70× and 0.80× (which admit 135,789 and 129,060 proteins, respectively) does not change which pathways are classified as universally present, universally absent, or variable.

The relaxed threshold was validated by four approaches. (1) Cross-referencing: of the 20,797 proteins admitted at 0.75× but not at 1.0×, 57% showed direct keyword agreement with their independent UniRef90 annotation (Supplementary Table 3), with the remainder reflecting nomenclature differences rather than functional disagreement. (2) *E. coli* negative control: applying the 0.75× pipeline to the *E. coli* K-12 MG1655 proteome (4,300 proteins) added 123 assignments (2.9%) beyond the 3,286 annotated at the strict threshold. Because the strict thresholds are already well-calibrated for *E. coli*, these additional assignments represent the noise floor of the relaxed cutoff — a modest rate that is tolerable given the substantial recovery of genuine *Pelagibacter* annotations. (3) Divergent sequence control: none of the 1,107 singleton proteins with no UniRef90 hit received a KO assignment at 0.75×. (4) Threshold sensitivity: the agreement rate between KofamScan and UniRef90 is stable across thresholds (57% at 0.75×, 56% at 0.70×, 60% at 0.80×), indicating that the quality of new assignments does not degrade as the threshold is relaxed within this range.

### Structural annotation recovers function where sequence methods fail

To complement sequence-based annotation, we predicted structures for 26,941 proteins using ESMFold and searched them against the AlphaFold Database using Foldseek. Of the 26,941 predicted structures, 22,216 (82.5%) matched an AlphaFold entry (e-value ≤ 10⁻³). Among the 5,980 folded proteins that were either annotated as hypothetical/DUF by UniRef90 or had no UniRef90 hit at all, 3,125 (52.3%) gained structural matches — functional information that sequence similarity alone could not provide. A total of 1,222 proteins had no detectable match by either sequence (UniRef90) or structure (Foldseek against AlphaFold DB). These are predominantly short (median 61 aa), singletons (77%), with depressed GC content (24.0% vs 29.1% genome mean), consistent with small HGT-acquired ORFs too short for reliable annotation by any current method. Twenty-one of the 1,222, however, are present in more than 10 genomes — conserved proteins of unknown function that may warrant experimental characterization. The most widespread (47 aa, present in 90 of 135 genomes across all three sources) occurs in a conserved operonic context alongside uracil-DNA glycosylase, a histidine phosphatase family protein, and a YbgC/FadM family acyl-CoA thioesterase — a neighborhood linking DNA repair and lipid metabolism. Its conservation across two-thirds of the genus, independent prediction by both Pyrodigal and NCBI’s PGAP pipeline, and fixed genomic context argue that it encodes a functional peptide, possibly regulatory, that has escaped detection by all current annotation methods. Of the 45 genomes lacking this protein, 44 also lack the entire operonic block — the uracil-DNA glycosylase, histidine phosphatase, and thioesterase are all absent together. Only one genome retains the block without the 47 aa peptide, carrying a clean deletion at its expected position. This co-segregation indicates that the peptide is an integral component of the operon, not an independent insertion, and illustrates the kind of fine-grained contextual analysis that is only possible with complete genomes where gene order is unambiguous.

### Auxotrophies are phylogenetically structured, not universal

*Pelagibacter* genomes lack biosynthetic pathways for several metabolites, creating obligate dependencies (auxotrophies) on exogenous nutrients. Previous work on a small number of isolates and SAGs documented auxotrophies for glycine, methionine, and several B vitamins (Tripp et al., 2008, 2009; Carini et al., 2013, 2014). However, it has remained unclear whether these auxotrophies are universal features of the genus or vary across lineages. Our dataset of 135 complete genomes allows, for the first time, a systematic assessment of pathway completeness at genus-wide scale where every apparent gene absence is genuine.

As a positive control, we verified that our pathway scoring correctly recapitulates the experimentally determined auxotrophies of *P. ubique* HTCC1062 (GCF_000012345), which is included in the dataset. Our scoring correctly identifies HTCC1062 as lacking assimilatory sulfate reduction, biotin biosynthesis, and de novo thiamine synthesis (while retaining thiamine salvage kinase) — all consistent with published culture experiments (Tripp et al., 2008, 2009; Carini et al., 2013, 2014). HTCC1062 also lacks threonine deaminase (K01754) by our threshold, predicting isoleucine auxotrophy; this has not been experimentally tested but is consistent with the defined medium used by Carini et al. (2013), which includes amino acids. Retained pathways (arginine, leucine, methionine salvage) are also correctly identified as present.

Module completeness analysis across 35 KEGG biosynthetic modules (Figure 7) reveals three categories of metabolic capability:

**Figure 7:**
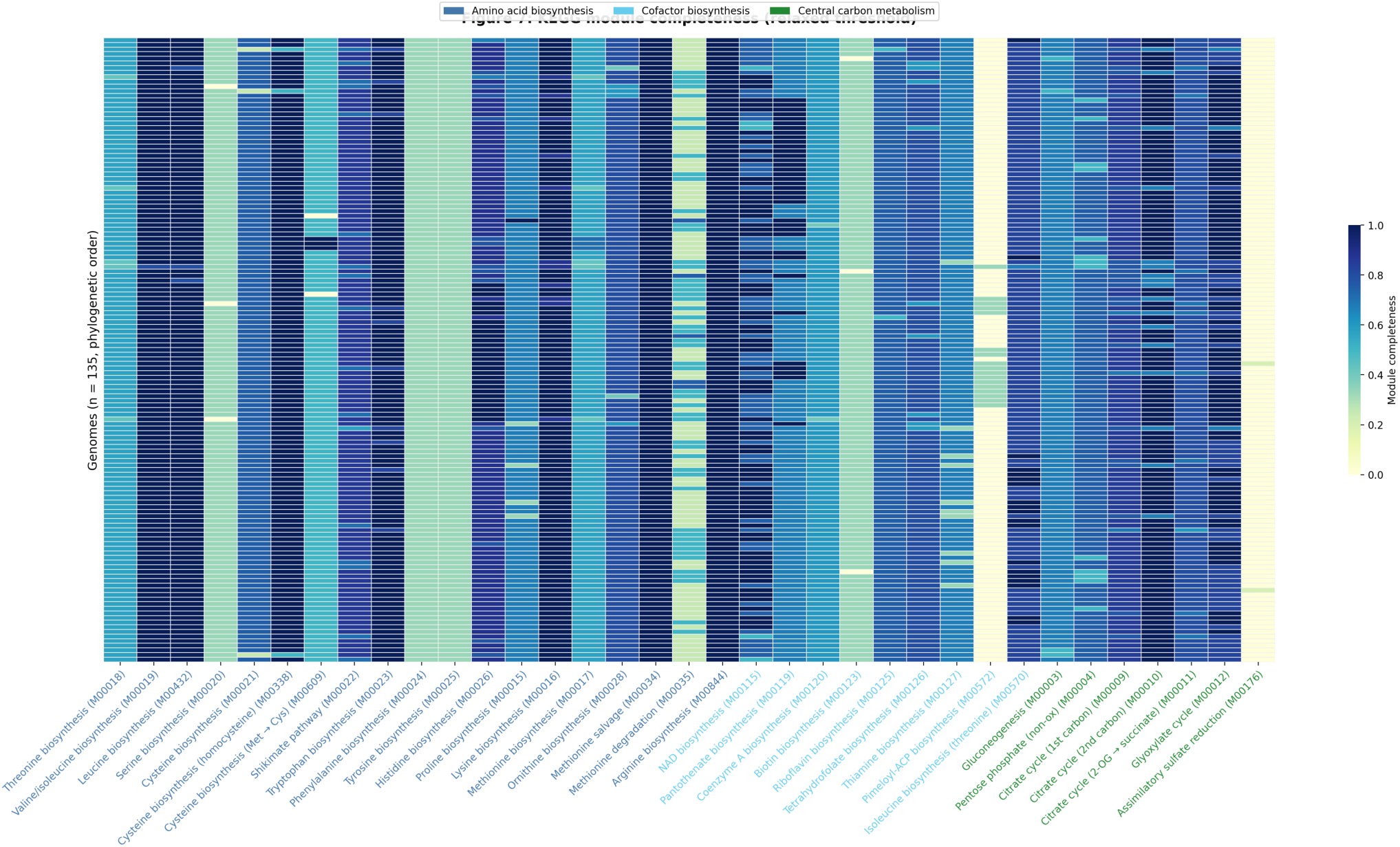
KEGG module completeness heatmap organized by phylogeny (relaxed threshold).

#### Universally retained pathways

Arginine biosynthesis is complete or nearly complete in most genomes: 126 retain all steps at the relaxed threshold, while 3 lack argD (K00818), 3 lack ornithine carbamoyltransferase (OTC, K00611), and 3 lack argE (K01438). The argE loss is compensated by the cyclic ornithine pathway via argJ (K00620, present in all 135 genomes), but the OTC and argD absences represent genuine pathway gaps. Methionine salvage, valine, leucine, and lysine biosynthesis are complete or nearly complete (≥98%) across all genomes. The TCA cycle, when assessed with the relaxed threshold that recovers the divergent citrate synthase and aconitase, is ≥83% complete in all genomes. These pathways are evidently essential under all ecological conditions experienced by *Pelagibacter*, or were present in the ancestor and have been maintained by selection.

#### Universally absent pathways

Assimilatory sulfate reduction is absent — no genome encodes a functional SO₄²⁻ → H₂S pathway. This represents an absolute, genus-wide dependency on reduced sulfur compounds (DMSP, methionine, cysteine). Biotin biosynthesis is also absent: the first committed step (8-amino-7-oxononanoate synthase, BioF/K00652) and the final step (biotin synthase, BioB/K01012) are genuinely missing, precluding de novo biotin synthesis. One intermediate enzyme (DAPA aminotransferase, BioA/K00833) is retained in 132 of 135 genomes at the relaxed threshold. BioA converts 7-keto-8-aminopelargonic acid (KAPA) to 7,8-diaminopelargonic acid (DAPA), an intermediate that cannot be further processed to biotin in the absence of BioD and BioB. Its near-universal retention may reflect a moonlighting function unrelated to biotin synthesis. The alternative biotin precursor pathway via BioC/BioH is also incomplete: BioC (K02169) is absent from all genomes and BioH (K02170) is present in only 17, ruling out both the canonical and alternative routes. Serine biosynthesis via the phosphorylated pathway (serA/serB/serC) is absent or incomplete in 129 of 135 genomes (the remaining 6 have only partial pathways), precluding de novo serine synthesis in the vast majority of species. Serine hydroxymethyltransferase GlyA (K00600) is present in all 135 genomes; this reversible enzyme interconverts serine and glycine while transferring a C1 unit to tetrahydrofolate, a reaction central to one-carbon metabolism (Sun et al., 2011). Culture experiments have established that *P. ubique* requires exogenous glycine for growth (Tripp et al., 2009; Carini et al., 2013). Threonine aldolase (K01620) is present in 120 of 135 genomes and can produce glycine from threonine, but this route depends on threonine availability. GlyA interconverts glycine and serine but cannot synthesize either de novo. Culture experiments confirm that *P. ubique* requires exogenous glycine for growth (Tripp et al., 2009; Carini et al., 2013), indicating that threonine aldolase does not provide sufficient glycine under laboratory conditions. GlyA generates serine from glycine as needed for protein synthesis and other metabolic demands. This glycine/serine dependency is a genus-wide feature. Phenylalanine and tyrosine biosynthesis are uniformly incomplete (33%), with chorismate mutase and aromatic aminotransferase absent. Whether these genus-wide absences reflect ancestral gene loss or reflect capabilities that were never present in the *Pelagibacter* lineage cannot be determined from our data alone; what is clear is that all 135 genomes share these dependencies regardless of species identity or habitat.

#### Variable pathways — phylogenetically structured variation

Several pathways show a pattern that is neither universally present nor universally absent, but varies across genomes:

- *Isoleucine*: threonine deaminase (K01754, the first committed step) is present in only 19 of 135 genomes. The remaining 116 genomes retain all downstream steps (shared with valine biosynthesis) but cannot initiate the isoleucine-specific branch.
- *Pantothenate*: ketopantoate reductase (K00077) is present in 36 genomes and absent in 99. Genomes without this enzyme cannot complete the pathway to vitamin B5.
- *NAD*: NAD⁺ synthase (K01916) is present in 74 genomes at the relaxed threshold. The remaining 61 genomes can synthesize nicotinate mononucleotide but cannot complete the final step — amidation of nicotinic acid adenine dinucleotide (NaAD) to NAD⁺.
- *Histidine*: histidinol-phosphatase (K01523) is present in only 29 genomes. Most genomes retain 7–8 of 9 histidine biosynthesis steps.
- *Glyoxylate cycle*: isocitrate lyase (K01637) is present in 99 genomes and absent in 36, determining whether the glyoxylate bypass is available.
- *Thiamine*: thiamine phosphate synthase (K03147) is absent in all genomes, but thiamine kinase (K00941) is present in 134 and thiamine-phosphate pyrophosphorylase (K00788) in 126 at the relaxed threshold. Thiamine salvage is thus near-universal, although de novo synthesis is not possible.

These variable pathways are the key finding. Parsimony-based permutation tests (10,000 tip shuffles, Bonferroni-corrected for 5 tests) confirm that the distribution of each variable pathway step on the phylogeny is significantly more clustered than expected by chance: histidine (2 changes observed vs 26.6 expected, p_adj < 0.0005), pantothenate (7 vs 31.9, p_adj < 0.0005), isoleucine (10 vs 18.1, p_adj < 0.0005), glyoxylate (13 vs 32.0, p_adj < 0.0005), and NAD (37 vs 44.6, p_adj = 0.056, borderline after correction). The phylogenetic distribution of these variable steps (Figure 8; Table 1; Supplementary Table 4) shows coherent patterns — losses and retentions cluster on the phylogeny rather than being randomly scattered — consistent with ecological specialization rather than neutral decay.

**Figure 8:**
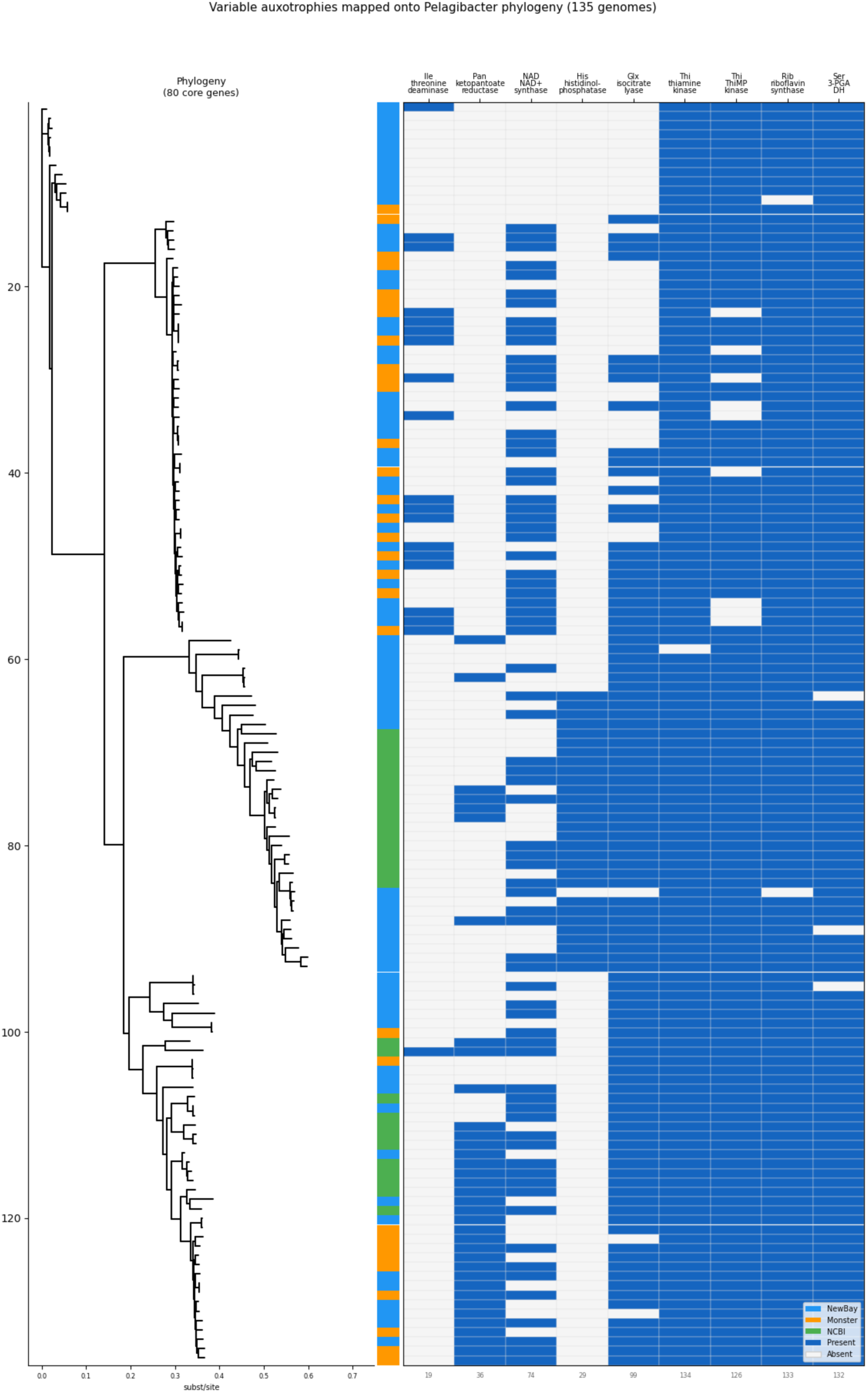
Auxotrophy heatmap. Rows = genomes ordered by phylogenomic tree. Columns = variable metabolic pathway steps. Cells colored by presence/absence. Clade annotations on left margin.

**Table 1:** Summary of variable pathway distributions across 135 genomes.

| Pathway | Key enzyme (KO) | Present | Absent | Parsimony changes | p_adj |
| --- | --- | --- | --- | --- | --- |
| Isoleucine | Threonine deaminase (K01754) | 19 | 116 | 10 vs 18.1 expected | <0.0005 |
| Pantothenate | Ketopantoate reductase (K00077) | 36 | 99 | 7 vs 31.9 | <0.0005 |
| NAD | NAD<sup>+</sup> synthase (K01916) | 74 | 61 | 37 vs 44.6 | 0.056 |
| Histidine | Histidinol-phosphatase (K01523) | 29 | 106 | 2 vs 26.6 | <0.0005 |
| Glyoxylate | Isocitrate lyase (K01637) | 99 | 36 | 13 vs 32.0 | <0.0005 |

### Transporters are constitutive, not compensatory

A natural prediction from variable auxotrophies is that genomes lacking a biosynthetic pathway should compensate with increased transporter capacity for the corresponding metabolite. We tested this by inventorying all transport systems across 135 genomes using the KofamScan KO assignments (relaxed threshold).

*Pelagibacter* genomes encode a consistent set of transport systems (Figure 9). TRAP transporters — the signature high-affinity uptake system of the lineage — are universal, with 2–9 periplasmic substrate-binding proteins (SBPs) per genome (mean 3.8). ABC transporters for polar amino acids (3–8 permease copies), branched-chain amino acids (2–8 copies), iron(III) (1–4 copies), and glycine betaine/proline (near-universal) are present in essentially every genome. DMSP demethylase (dmdA) is universal with 1–3 copies, confirming that all *Pelagibacter* species can demethylate this key organosulfur compound. Additionally, 27 of 135 genomes carry DMSP lyase DddP (K16953), which cleaves DMSP to release dimethyl sulfide (DMS) and acrylate. However, Sun et al. (2016) demonstrated that *P. ubique* HTCC1062 produces both DMS and methanethiol from DMSP simultaneously despite lacking DddP, implying the existence of additional, uncharacterized DMSP cleavage enzymes. Whether the 108 genomes without DddP retain DMS-producing capacity through other DMSP lyases or as-yet-unidentified enzymes cannot be determined from genomic data alone. This variable distribution of DMSP processing routes represents another axis of metabolic differentiation across the genus, with potential implications for marine sulfur cycling (Sun et al., 2016). Spermidine/putrescine permeases and Na⁺/H⁺ antiporters are also universal.

**Figure 9:**
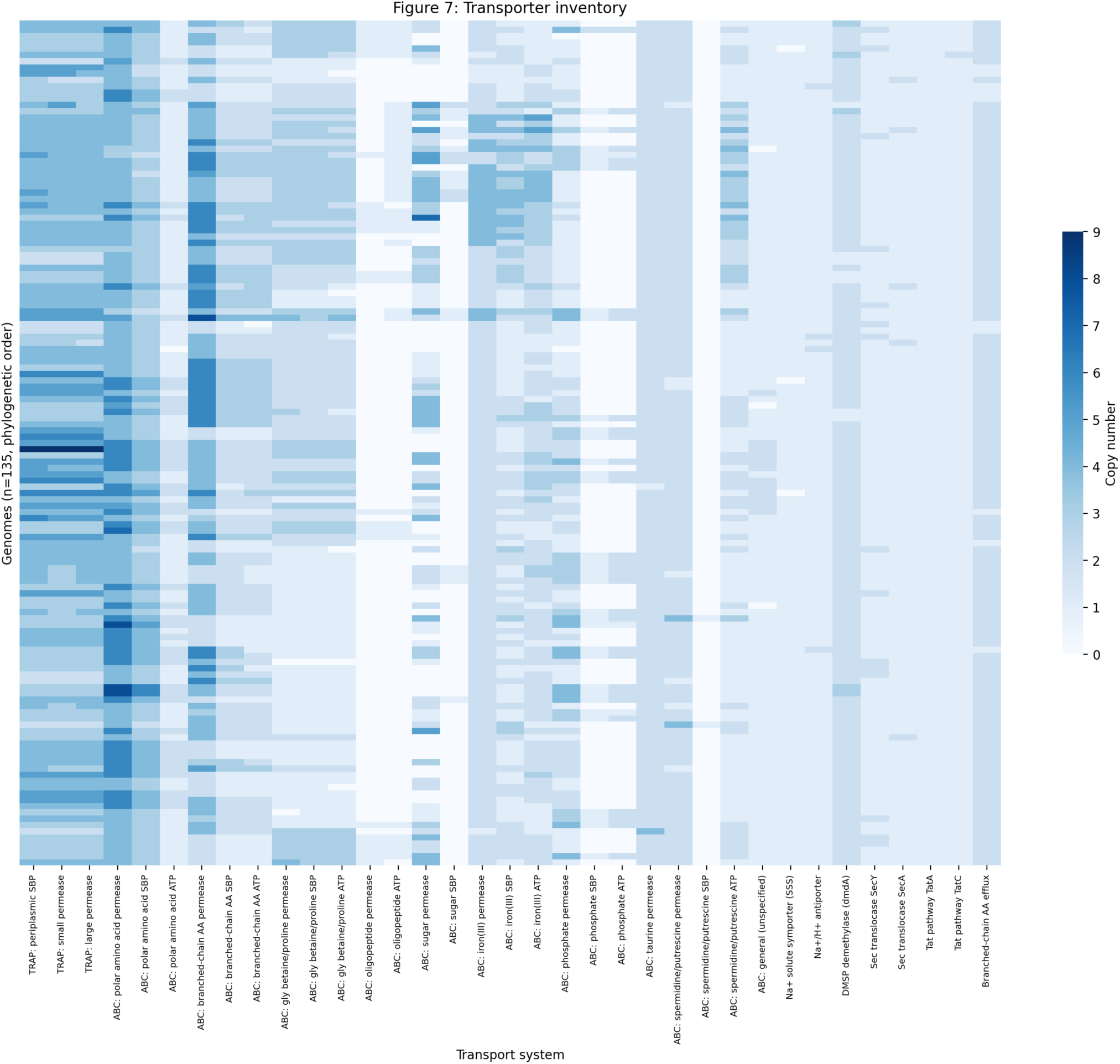
Transporter inventory heatmap. Genomes ordered by phylogeny, columns = transport systems, colored by copy number.

The transport complement is notably uniform despite the variable biosynthetic capabilities documented above. We tested whether genomes lacking a biosynthetic pathway compensate with additional substrate-specific transporters, comparing copy numbers for each variable pathway’s corresponding transporter (Mann-Whitney U test, Bonferroni-corrected for 3 tests). Genomes lacking isoleucine biosynthesis carry the same number of branched-chain amino acid SBPs as those with the pathway (2.0 vs 2.1; U = 1147, p = 0.76). Genomes lacking histidine biosynthesis show no significant difference in polar amino acid SBPs (3.3 vs 3.5; U = 1876, p_adj = 0.11). Genomes lacking serine biosynthesis have identical polar amino acid transporter counts to the few with partial serine pathways (3.3 copies each; U = 193, p = 0.94). No comparison was significant after correction. This absence of compensation implies that the transport systems are constitutive: they are maintained at a fixed copy number regardless of biosynthetic capability, presumably because exogenous nutrient uptake is always advantageous in the oligotrophic marine environment, whether or not the cell can synthesize the metabolite itself.

This uniformity reframes the variable auxotrophies. All *Pelagibacter* genomes import the same metabolites at the same capacity — transporters do not compensate for biosynthetic loss because they were never contingent on it. Import is constitutive; biosynthesis is the variable. A genome that retains a biosynthetic pathway can survive when the exogenous metabolite is scarce; one that has lost it cannot, regardless of transporter copy number. The cost of maintaining biosynthesis — genes, expression, precursors — only pays off when supply is unreliable. The four phylogenetic groups visible in Figure 8, each with a distinct combination of retained and lost pathways, may therefore reflect adaptation to environments that differ in which metabolites are reliably available.

### Gene order is not conserved across species

Previous analyses of small numbers of SAR11 genomes reported high synteny conservation: Wilhelm et al. (2007) found 96% synteny among a handful of genomes from the Sargasso Sea, and Grote et al. (2012) described SAR11 as exhibiting “high synteny” with rearrangement confined to hypervariable regions flanked by rRNA operons. With only 2–5 genomes from closely related lineages available at the time, this conclusion was reasonable but reflected limited sampling.

Our analysis of gene order across 135 complete genomes reveals a different picture. We defined synteny in terms of gene adjacencies — consecutive gene pairs (by cluster assignment and strand) in positional order along each chromosome. Of 40,011 unique adjacencies observed across all genomes, only 2 were universal (present in all 135 genomes) and only 41 were near-universal (present in ≥130). In contrast, 25,981 adjacencies (65%) were singletons found in a single genome. The adjacency frequency spectrum mirrors the gene frequency spectrum: a tiny conserved core and a vast cloud of genome-specific arrangements (Figure 10A).

**Figure 10:**
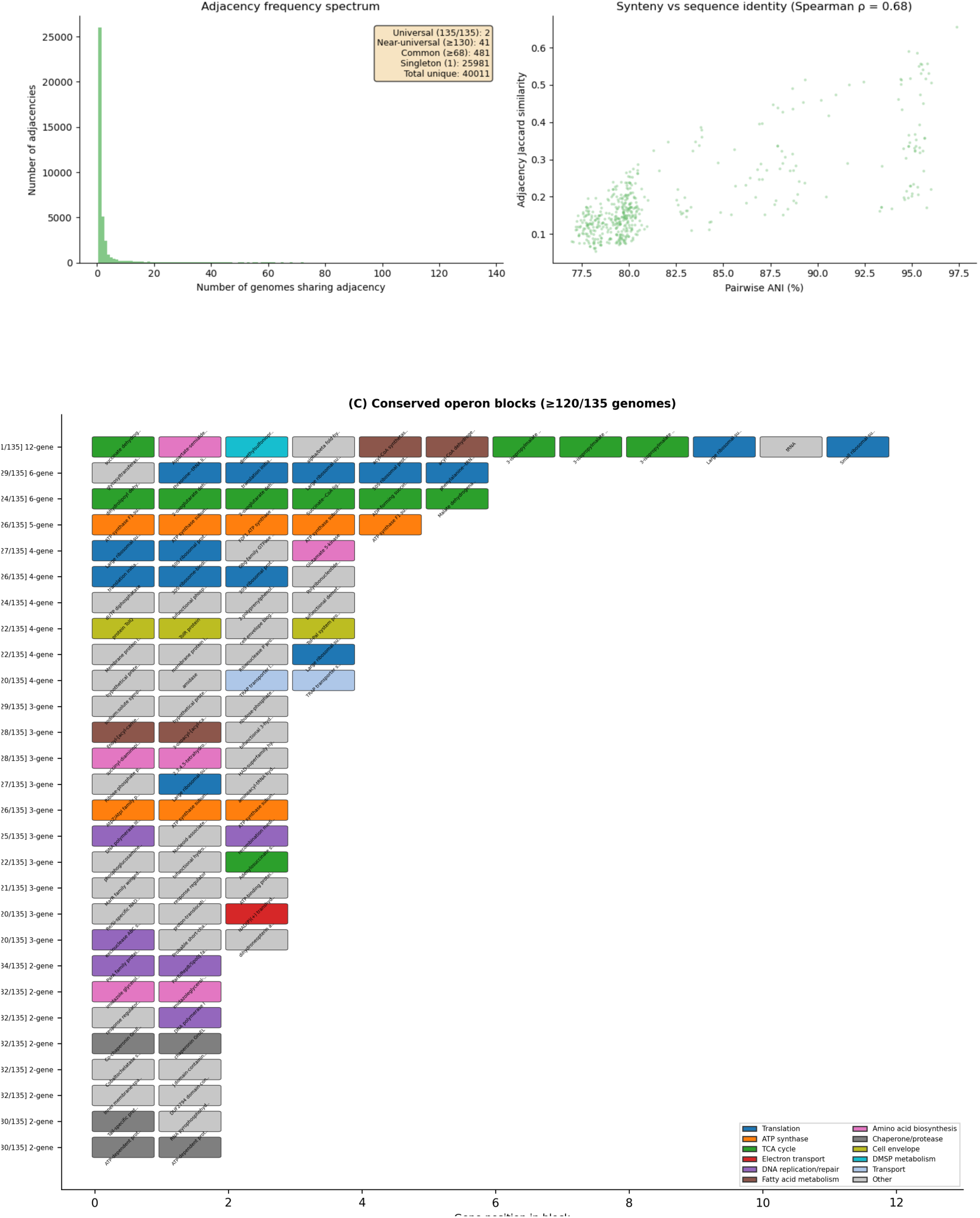
(A) Adjacency frequency spectrum. (B) Pairwise synteny Jaccard vs ANI scatterplot. (C) Conserved operon block diagram.

Pairwise synteny conservation correlates strongly with sequence identity but decays rapidly (Figure 10B). Within-species genome pairs (>95% ANI) share ∼75–82% of gene adjacencies (Jaccard similarity), confirming that closely related genomes do maintain high synteny as previously reported. However, between-species pairs show rapid loss: at ∼85% ANI, Jaccard similarity drops to ∼10%, and the most divergent pairs (76–80% ANI) share fewer than 9% of adjacencies. The mean pairwise Jaccard across 500 random genome pairs was 0.19 — far from the near-universal conservation implied by earlier studies.

This finding has implications for understanding metagenome assembly of *Pelagibacter*. The primary assembly barrier is high nucleotide identity in conserved genes shared between co-occurring species — stretches where the assembler cannot distinguish one genome from another. The HVR compounds this problem: because it evolves more rapidly than the rest of the genome, co-occurring species that are near-identical across their conserved backbone carry completely different HVR content flanked by the same shared sequence on both sides. The assembler sees the shared flanks, attempts to merge the genomes, and encounters divergent HVR paths that it cannot assign to individual genomes without sufficient read coverage spanning the transitions. The scrambled gene order between species creates additional species-specific signatures that in principle aid assembly, but resolving these paths still requires reads that span the shared-to-unique boundaries, which explains why increased sequencing depth recovers additional species (see below).

To characterize this conserved core more systematically, we predicted operons across all 135 genomes using intergenic distance (<100 bp) and co-directionality as criteria. On average, 87% of genes in each genome fall within a predicted operon (mean 272 operons per genome, mean size 4.6 genes). We then identified operon adjacencies conserved across genomes — gene pairs that remain co-transcribed despite the extensive inter-operon rearrangement.

Five operon adjacencies were universal (present in all 135 genomes): FabB → FabA (fatty acid biosynthesis), bL21 → L27 and S17 → L29 (ribosomal proteins), cytochrome b → cytochrome c1 (respiratory electron transport), and an ABC transporter permease → ATP-binding protein pair. Three of these were not detected as universal in the whole-genome synteny analysis above because the operon analysis uses a relaxed definition: gene pairs need only be co-directional and within 100 bp, regardless of whether intervening unclustered genes disrupt the strict positional adjacency. An additional 89 adjacencies were near-universal (≥130/135).

Annotation of the 153 unique gene clusters involved in these 94 conserved adjacencies (via UniRef90) revealed distinct functional categories:

- **Ribosome and translation** (31 adjacencies): the largest conserved block. Multiple long chains of ribosomal proteins remain linked, including L24 → uL14 → S17 → L29 → L16, L1 → L11 → NusG (transcription termination), and S19 → L22. Translation factors (IF-3, EF-Tu) and aminoacyl-tRNA synthetases are also conserved alongside ribosomal protein genes.
- **TCA cycle and energy metabolism** (8): succinate-CoA ligase alpha → beta → malate dehydrogenase; electron transfer flavoprotein beta → alpha subunits.
- **Amino acid biosynthesis** (8): leucine (LeuB → LeuD → LeuC), tryptophan (anthranilate phosphoribosyltransferase → anthranilate synthase), histidine (HisH → HisB, HisF → HisB).
- **DNA replication and repair** (6): including DNA polymerase III subunits and gyrase.
- **ATP synthase** (5): F1 subunits (gamma → alpha → delta) and F0 subunits (a → c) linked in separate conserved blocks.
- **Respiratory chain** (4+): NADH dehydrogenase (Complex I) subunits J → K, B → A, and B → C remain linked; cytochrome c oxidase assembly protein → subunit 3.
- **Fatty acid metabolism** (4): FabB → FabA (universal), plus acyl-CoA synthetase → dehydrogenase.
- **Chaperone and protease systems** (4): GroES → GroEL, ClpX → ClpP, HslV → HslU.
- **Iron-sulfur cluster assembly** (3): the SUF system, with aminotransferase → SufU → SufS linked.
- **Chromosome partitioning** (2): ParA → ParB.
- **Molybdenum cofactor biosynthesis** (2): MoaC → MoaA.
- **Sulfite oxidation** (2): dissimilatory adenylylsulfate reductase subunits (AprA → AprB).

Chaining conserved pairwise adjacencies into maximal blocks revealed 58 multi-gene operon units conserved across ≥120 of 135 genomes (Figure 10C; Supplementary Table 5). The largest is a 12-gene conserved gene block spanning succinate dehydrogenase, aspartate-semialdehyde dehydrogenase, DMSP demethylase, acyl-CoA metabolism, three leucine biosynthesis enzymes (LeuBDC), and ribosomal proteins bL19 and bS16 — a chain linking energy metabolism, amino acid biosynthesis, and translation that is maintained in 121 or more genomes. Other notable conserved blocks include a 6-gene TCA cycle operon (dihydrolipoyl dehydrogenase through malate dehydrogenase), the 5-gene F1-ATP synthase operon (epsilon → beta → gamma → alpha → delta), a 6-gene block linking a glycosyltransferase to five translation components (threonine-tRNA ligase → IF-3 → bL35 → L20 → phenylalanine-tRNA ligase), the TolQRAB operon (4 of 5 Tol-Pal system components; TolQ → TolR → TolA → TolB), and a 3-gene fatty acid synthesis block (FabI → FabB → FabA).

The picture that emerges is of a genome organized as a set of conserved operonic blocks — multi-gene units encoding multi-subunit complexes and sequential pathway enzymes — that are extensively shuffled relative to each other across species. The operonic blocks are under strong selection to maintain co-transcription, while the order in which they appear along the chromosome is free to vary. Rearrangement occurs at the inter-operon level; the interior of each operon remains intact.

### Sequencing depth affects *Pelagibacter* species recovery

A unique feature of our dataset is the inclusion of two assemblies from the same sample: station 8 summer (8S), sequenced with 2 ONT flow cells (standard depth), and 0S, the same sample with 4 additional flow cells added to the original 2 (6 total, approximately 3× depth). Because 0S is a superset of 8S, any species recovered in 0S but not 8S is attributable to the additional sequencing depth. This controlled comparison directly addresses the relationship between sequencing effort and *Pelagibacter* diversity recovery.

At standard depth (8S), 20 complete genomes spanning 4 species were recovered (Figure 11). At elevated depth (0S), 31 complete genomes spanning 9 species were recovered. Of the 9 species in 0S, 3 were shared with 8S, 3 were found at other standard-depth stations but not at 8S, and 3 were not found in any of the 16 standard-depth assemblies. One species present at 8S (*P.* sp001438335, a single genome) was not recovered in 0S, likely because the additional coverage from closely related species complicated the assembly graph for this low-abundance genome. We searched for these 3 depth-dependent species in the standard-depth assemblies from all samples using skani. One species (M_u12081019) was absent from all standard-depth assemblies entirely; the other two were present as high-ANI fragments — partial genomic sequences with >99% nucleotide identity to the 0S reference, but failing to assemble into complete contigs.

**Figure 11:**
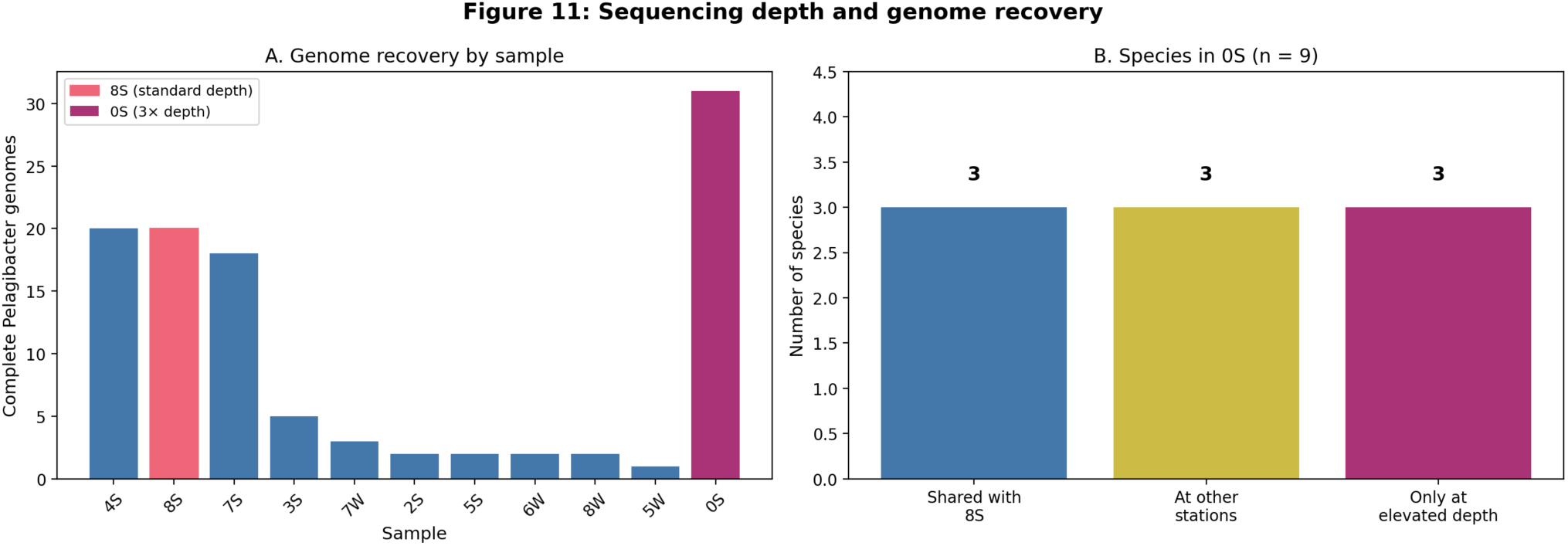
(A) Genome recovery by sample — 8S highlighted, 0S (3× depth) shown separately. (B) Species in 0S: 3 shared with 8S, 3 at other stations, 3 unique to elevated depth.

These results demonstrate that standard metagenomic sequencing depth systematically underestimates *Pelagibacter* diversity. The mechanism is not absence of the organisms but failure of assembly: because co-occurring species share near-identical backbone sequence but carry different HVR content, the assembler must decompose pooled coverage from all species in the flanking regions to resolve individual genome paths through the HVR. Each species contributes only a fraction of the total coverage at these shared loci, so per-species depth is the limiting factor. The 3× increase in sequencing depth at 0S more than doubled the species count at this station (9 vs 4) and recovered 3 species that were not assembled at standard depth anywhere in the transect. This comparison is unreplicated (n = 1 station), so the specific recovery rate should not be generalized, but the qualitative finding — that increased depth recovers additional species present as fragments at standard depth — is consistent across the depth-dependent species examined. The true diversity at this station is likely higher still.

## Discussion

### *Pelagibacter* diversity is vastly undersampled

Our recovery of 44 novel species from a single estuary — representing 85% of the 52 species in our combined dataset — illustrates how much *Pelagibacter* diversity remains undescribed. The GTDB currently contains hundreds of *Pelagibacter* species-level taxa, but these are overwhelmingly represented by fragmentary MAGs and SAGs. Complete genomes, which are necessary for unambiguous assessment of gene content and pathway completeness, remain scarce. The addition of 106 new complete genomes from our study (75 SFE + 31 deeply sequenced 0S) more than quadruples the number of closed *Pelagibacter* genomes available in public databases.

This novelty rate is notable given that the SFE is not an underexplored environment. Previous metagenomic studies have characterized its microbial communities using both short-read (Rasmussen & Francis, 2022, 2023) and long-read (Lui & Nielsen, 2024) approaches. Brackish-water relatives of *Pelagibacter* (SAR11 subclade IIIa) have been specifically studied in San Francisco Bay using targeted cultivation and MAG recovery (Lanclos et al., 2023). Yet more than half the *Pelagibacter* species we recovered have no close relative in any reference database. This suggests that the gap in described diversity is not specific to understudied environments but reflects a fundamental limitation in the methods — particularly short-read metagenomics and cultivation — that have been used to characterize *Pelagibacter* populations.

The species accumulation curve (Figure 2B) does not plateau, indicating that even our 135 genomes do not capture the full *Pelagibacter* diversity present in the SFE. This is consistent with the observation that the deeply sequenced 0S station yielded 3 species not found in any standard-depth assembly and more than doubled the species count at that station. Together, these results indicate that systematic, deep, long-read metagenomic sequencing of additional environments will continue to reveal new *Pelagibacter* species for the foreseeable future.

A search of all 1,442 genomes classified as *Pelagibacter* at NCBI (excluding the 31 already in our dataset) against our 135 genomes using skani quantifies both the utility of our collection and the scope of remaining diversity. Of 1,442 NCBI genomes, 1,132 (79%) share detectable ANI (>80%) with at least one of ours; 420 (29%) fall within our species at ≥95% ANI, providing the first complete reference for 331 contig-level MAGs and 60 scaffold-level assemblies that previously lacked a closed genome in their species. However, 1,022 NCBI genomes (71%) represent species not covered by our 52. The 310 genomes below ANI detection likely include non-*Pelagibacter* SAR11 lineages (e.g., IMCC9063 subclade II, *Fonsibacter*) that NCBI classifies under the broader “Candidatus Pelagibacter” taxonomy. Our 135 genomes thus serve as complete references for a substantial fraction of existing public data while covering only a fraction of the genus-level diversity — further underscoring the scale of undescribed *Pelagibacter* species.

### Metabolic dependencies are structured, not uniform

The standard interpretation of *Pelagibacter*’s compact genome holds that auxotrophies reflect the loss of dispensable biosynthetic pathways — a natural consequence of genome streamlining in organisms that can obtain these metabolites from the environment (Giovannoni et al., 2005; Giovannoni, 2017). Under this model, auxotrophies should be widespread and relatively uniform across the genus, accumulating stochastically as lineages shed unnecessary genes.

Our survey of 35 biosynthetic modules across 135 complete genomes reveals a more structured picture. Metabolic dependencies fall into three categories that differ in their evolutionary implications.

The first category — genus-wide dependencies — includes biotin biosynthesis (absent in all 135 genomes), assimilatory sulfate reduction (absent in all), and glycine/serine (serine biosynthesis absent in 129 of 135; the reversible enzyme GlyA is present in all genomes but cannot synthesize either amino acid de novo). These are near-universal or universal commitments to exogenous supply. All or nearly all *Pelagibacter* species depend on external biotin, reduced sulfur compounds, and glycine. Whether these dependencies arose through ancestral gene loss or were never acquired by the *Pelagibacter* lineage cannot be determined from our data; what is clear is that they are fixed features of the genus.

The second category — universally retained pathways — includes arginine, valine, leucine, lysine biosynthesis, and the TCA cycle. These are maintained in all or nearly all genomes, implying that they are essential regardless of ecological context. Their universal retention constrains the minimum metabolic independence of the genus.

The third category — variable pathways — is the key finding. Isoleucine biosynthesis (threonine deaminase present in 19 of 135 genomes), pantothenate (ketopantoate reductase in 36), NAD (NAD⁺ synthase in 74), histidine (histidinol-phosphatase in 29), and the glyoxylate cycle (isocitrate lyase in 99) each show a pattern of presence in some lineages and absence in others. These are not random losses scattered across the phylogeny — four of the five show significant phylogenetic clustering (histidine, pantothenate, isoleucine, and glyoxylate; p_adj < 0.0005), with NAD borderline (p_adj = 0.056). The presence/absence patterns (Figure 8) indicate that pathway gains or losses occurred at specific points in the *Pelagibacter* radiation and were subsequently inherited by descendant lineages.

The variable auxotrophies have direct ecological implications. Our transporter inventory shows that all *Pelagibacter* genomes maintain the same complement of amino acid and nutrient uptake systems regardless of biosynthetic capability. Genomes lacking isoleucine biosynthesis do not compensate with additional branched-chain amino acid transporters; genomes lacking serine biosynthesis have the same polar amino acid transporter copy number as those with the pathway. This constitutive transport architecture means that auxotrophic genomes are genuinely more dependent on exogenous metabolite supply — they use the same uptake capacity but have no biosynthetic fallback when the environmental supply is insufficient.

This creates a potential mechanism for niche partitioning. A *Pelagibacter* species that retains isoleucine biosynthesis can persist when exogenous isoleucine is scarce; one that has lost the pathway cannot. If different metabolites fluctuate independently in the environment — as they likely do across estuarine gradients, seasons, and depth strata — then different combinations of retained and lost pathways would adapt different species to different nutrient regimes. The variable auxotrophies would thus represent not the stochastic decay of a once-complete genome, but the diversification of metabolic strategies across a genus whose members partition a heterogeneous nutrient landscape.

The phylogenetic placement of *Pelagibacter* within the Alphaproteobacteria, whose members generally possess larger genomes and more complete biosynthetic repertoires, is consistent with some degree of genome reduction having occurred in the *Pelagibacter* lineage. However, the magnitude and timing of this reduction remain uncertain, and the possibility that the *Pelagibacter* ancestor possessed a genome closer in size to the extant genus than to its larger-genomed relatives cannot be excluded. What our data demonstrate is that, regardless of the ancestral state, the *pattern* of metabolic loss is structured rather than uniform — different lineages have arrived at different metabolic configurations. This is the novel contribution: not whether reduction occurred (which the phylogenetic context suggests it did, at least to some degree) but that the process was ecologically differentiated, producing distinct metabolic strategies across a genus that partitions a heterogeneous nutrient landscape.

### Complete genomes vs. MAGs and SAGs

Our dataset highlights a critical advantage of complete genomes over MAGs and SAGs for comparative genomics of *Pelagibacter*. With genomes of ∼1.3 Mbp and coding densities approaching 96%, there is very little non-coding space. In such compact genomes, the loss of even a few genes — whether through genuine biological deletion or through assembly/ binning artifact — can alter the apparent metabolic profile of a genome. A single missing gene can determine whether a biosynthetic pathway is scored as complete or absent.

MAGs, by definition, are fragmented and potentially contaminated assemblies reconstructed through binning of metagenomic contigs. For *Pelagibacter*, the challenges are compounded: (1) the high sequence similarity between co-occurring species means that binning algorithms may merge contigs from different species, (2) the conserved genomic regions that cause assembly fragmentation also create chimeric contigs that span species boundaries, and (3) the small genome size means that even modest contamination can inflate apparent gene content by >10%.

SAGs avoid the contamination and chimera problems of MAGs but suffer from amplification bias, resulting in incomplete genome coverage. For *Pelagibacter*, typical SAG completeness values are 30–70%, making it impossible to determine whether a pathway gene is genuinely absent or simply not captured.

The 135 complete genomes in our dataset eliminate these ambiguities. Every gene present in each genome is accounted for, and every apparent absence is genuine. This is essential for the auxotrophy analysis (Figure 8), where the distinction between “gene absent” and “gene not recovered” is the difference between a meaningful biological conclusion and an artifact.

### Methodological insight: sequencing depth and the *Pelagibacter* assembly bottleneck

Our controlled comparison of sequencing depths provides direct evidence that *Pelagibacter* diversity is systematically underestimated by standard metagenomic approaches. The key insight is that the bottleneck is not detection but assembly: organisms are present in the sample and their reads are present in the data, but the assembly algorithm cannot resolve their genomes because conserved regions shared with other co-occurring species create ambiguities in the assembly graph.

This has practical implications for study design. For projects aiming to catalog the full *Pelagibacter* diversity at a site, our results suggest that standard metagenomic sequencing depths (e.g., 2 ONT flow cells per sample) are insufficient. The recovery of 3 additional species at 3× depth — and the doubling of the species count from 4 to 9 — indicates substantial but not exhausted returns, and further increases in depth — or complementary approaches such as read-cloud or linked-read sequencing — may be needed to approach saturation.

For the most abundant bacterial genus in the ocean, we are still discovering new species with each increase in methodological resolution.

## Methods

### Sample collection and sequencing

Water samples were collected from 8 stations along the SFE salinity gradient during summer 2022 and winter 2023. Station locations and environmental metadata are available through the U.S. Geological Survey (USGS) California Water Science Center monitoring program. DNA extraction, library preparation, and Oxford Nanopore Technologies (ONT) long-read sequencing were performed as described in Lui & Nielsen (2024). Each sample was sequenced with 2 ONT flow cells. For station 8 summer, 4 additional flow cells were subsequently sequenced and combined with the original 2 to produce the 0S assembly (6 total flow cells, ∼3× depth) used in the depth comparison analysis.

### Metagenome assembly

All assemblies were performed with myloasm v0.4.0 (Shaw, Marin & Li, 2026), a long-read metagenome assembler designed for high-resolution strain resolution through polymorphic k-mer-based string graph construction. Default R10 nanopore parameters were used (compression ratio c = 11, minimum overlap 500 bp, quality value cutoff 90% identity). Circular contigs were identified from the myloasm output, which flags circularization when the assembler detects overlap between the start and end of a contig during graph traversal.

### *Pelagibacter* genome identification and quality assessment

Contigs ≥500 kbp were assessed for genome completeness and contamination using CheckM2 v1.1.0 (Chklovski et al., 2023) with the UniRef100.KO.1 DIAMOND database. Contigs with ≥90% estimated completeness and <5% estimated contamination were classified as high-quality (HQ) genomes. Taxonomic classification was performed with GTDB-Tk v2.6.1 `classify_wf` (Chaumeil et al., 2022) using GTDB release R226 (Parks et al., 2025). Genomes classified as *Pelagibacter* (g Pelagibacter) and assembled as complete single-contig genomes were selected for this study.

### NCBI reference genomes

All complete *Pelagibacter* genome assemblies were downloaded from NCBI (n = 31; 30 RefSeq + 1 GenBank-only). Two were excluded after quality and taxonomic assessment (see Results), yielding 29 NCBI genomes in the final dataset. [Accession numbers listed in Supplementary Table 6.]

### Genome reorientation

All 135 genomes were reoriented to begin at the dnaA gene using dnaapler v1.3.0 (Bouras et al., 2024) in bulk chromosome mode. This ensures consistent start positions for synteny comparisons and chromosomal position analyses.

### Gene prediction

Protein-coding genes were predicted using Pyrodigal v3.6.3 (Larralde, 2022) in single-genome mode (`-p single`), which trains gene-finding parameters individually on each genome. This is preferred over meta mode for complete genomes, particularly for *Pelagibacter*’s extreme AT-richness (∼29% GC), where per-genome training provides more accurate start-site prediction and gene boundary identification.

### tRNA prediction

tRNA genes were predicted using tRNAscan-SE v2.0.12 (Chan & Lowe, 2019) in bacterial mode (`-B`) with 4 threads per genome. The initiator methionine tRNA is reported by tRNAscan-SE as “fMet” and the elongator methionine tRNA as “Ile2” (CAT anticodon); both were counted as methionine tRNA for amino acid coverage assessment. Genomes with incomplete amino acid coverage were re-analyzed with tRNAscan-SE in maximum sensitivity mode (`--max`, bypassing the HMM pre-filter) and independently with ARAGORN v1.2.41 (Laslett & Canback, 2004) to resolve discrepancies.

### Average nucleotide identity (ANI)

Pairwise ANI was calculated using skani v0.3.1 (Shaw & Yu, 2023) in triangle mode with the `--slow` preset. Genome pairs below skani’s detection threshold (∼80% ANI) return no result and were treated as 0% ANI in the distance matrix. Species-level clusters were defined at a 95% ANI threshold using single-linkage connected components (NetworkX v3.3 in Python v3.12): a graph was constructed with genomes as nodes and edges between pairs sharing ≥95% ANI, and each connected component was treated as a species. A species accumulation curve was generated by randomly permuting the order of the 16 metagenomic samples (excluding 0S) over 50 iterations and recording the cumulative number of species observed at each step.

### Phylogenomic analysis

Single-copy core gene clusters were identified from the MMseqs2 pangenome clustering (see below) as clusters containing exactly one member per genome across all 135 genomes, yielding 80 clusters. For each cluster, protein sequences were extracted with genome names as identifiers and aligned using MAFFT v7.525 (Katoh & Standley, 2013) with the `--auto` strategy (which selected L-INS-i for these alignments) and 4 threads per alignment (8 alignments in parallel). No alignment trimming was applied, as the low gap content in these single-copy core genes from complete genomes (0% missing data) makes trimming unnecessary. The 80 alignments were concatenated into a supermatrix of 135 taxa and 17,509 aligned amino acid positions, with a corresponding partition file assigning the LG+G4+F substitution model to each gene partition. Maximum-likelihood phylogenetic inference was performed with IQ-TREE v3.0.1 (Wong et al., 2025; Minh et al., 2020) using the partitioned model, 64 threads, and 1000 ultrafast bootstrap replicates (UFBoot2; Hoang et al., 2018). The resulting tree (log-likelihood −300,590.8, total tree length 4.10) was used as the backbone for all subsequent analyses including auxotrophy mapping and pangenome ordering. A rooted tree was inferred by adding *Ca.* Pelagibacter sp. IMCC9063 (GCF_000195085; SAR11 subclade II, excluded from the main analysis due to extreme divergence) as an outgroup. IMCC9063 sequences were added to the 72 core gene alignments for which orthologs were found (mean 51% amino acid identity) using MAFFT `--add ––keeplength`, which inserts the new sequence without altering the existing alignment columns. For the 8 core genes lacking an IMCC9063 ortholog, the outgroup was represented by gaps. The rooted tree was inferred with the same IQ-TREE settings (136 taxa, 17,509 positions, 80 partitions, 1000 UFBoot replicates, outgroup specified with `-o GCF_000195085`).

### Expanded phylogeny

To assess whether the 135-genome collection captures the broader phylogenetic diversity of the genus, an expanded tree was constructed by adding 89 additional high-quality *Pelagibacter* genomes from NCBI (≥90% completeness, <5% contamination, confirmed as g Pelagibacter by GTDB-Tk). Proteins were predicted with Pyrodigal and searched against the 80 core cluster representatives using MMseqs2 (sensitivity 7.5, 70% identity, 80% coverage). Orthologs were added to the existing core gene alignments using MAFFT `--add ––keeplength`. The resulting 225-taxon supermatrix (17,509 positions, 80 LG+F+G4 partitions) was analyzed with IQ-TREE using the same settings as the 135-taxon tree (1000 UFBoot replicates, 32 threads). The tree was rooted externally using BioPython after IQ-TREE’s internal rooting step failed.

### Pangenome analysis

All predicted proteins from the 135 genomes (192,716 sequences) were clustered using MMseqs2 v18.8cc5c `easy-cluster` with minimum sequence identity of 70%, minimum bidirectional coverage of 80% (`--cov-mode 0`), set-cover clustering mode (`--cluster-mode 0`, the default), and default sensitivity (s = 4). A presence/absence matrix was constructed by mapping each protein to its source genome and cluster assignment. Multi-copy instances (paralogs within the same genome assigned to the same cluster) were recorded. Truncations were flagged when a protein was shorter than 80% of the median length within its cluster, as a potential indicator of assembly-induced frameshifts. Pangenome accumulation curves were generated from 50 random permutations of genome addition order (random seed 42); at each step, the number of total clusters, core clusters (present in all genomes added so far), and newly observed clusters was recorded. Gene content distance between genomes was computed as Jaccard distance on the binary presence/absence profiles and compared to ANI distance using a Mantel test (Spearman rank correlation on distance matrices with 9,999 permutations, implemented in Python v3.12 with SciPy v1.12.0; all 9,045 pairwise comparisons included). To test sensitivity to clustering parameters, the full analysis was repeated at 50% and 90% identity thresholds (with 80% coverage held constant); key metrics (total clusters, core count, singleton fraction, accumulation curve slope) were compared across the three thresholds.

### Assembly frameshift detection

Assembly-induced frameshifts were detected by two methods. First, split genes were identified as pairs of adjacent ORFs on the same strand whose protein sequences both match the same UniRef90 target (e-value ≤ 10⁻⁵) and whose combined length approximates the target length. Second, truncated-with-gap candidates were identified as proteins with query/target length ratio <0.7 (missing >50 amino acids) followed by an intergenic gap between 0.5× and 2.0× the expected size of the missing portion (missing amino acids × 3 bp). Normal intergenic distance in *Pelagibacter* is 4 bp (median), so gaps >200 bp following a truncated protein are highly anomalous and consistent with an uncalled gene fragment.

### Chromosomal position analysis

All genomes were reoriented to start at dnaA (see Genome reorientation above). For each protein-coding gene, its chromosomal position was expressed as a fraction of total genome length (0 = origin, 0.5 = terminus). Gene conservation was measured as the frequency of each gene’s cluster across all 135 genomes. The relationship between chromosomal position and conservation was assessed by comparing mean cluster frequency and core gene fraction in origin-proximal (position 0–0.25 and 0.75–1.0) versus terminus-proximal (position 0.25–0.75) regions.

### Functional annotation

All 192,716 predicted proteins were searched against UniRef90 (release 2024_02; UniProt Consortium, 2024) using MMseqs2 v18.8cc5c `easy-search` with an e-value threshold of 1 × 10⁻⁵, retaining the top hit per query. Output fields included query and target lengths, alignment length, percent identity, and target description. Proteins with query/target length ratios below 0.8 were flagged as potentially truncated for downstream quality assessment.

KEGG ortholog assignments were made using KofamScan v1.3.0 (Aramaki et al., 2020) with KEGG HMM profiles (downloaded March 2026) and the associated ko_list adaptive thresholds. All 192,716 proteins were searched in a single run using 32 CPUs. Because KofamScan’s profile-specific score thresholds are calibrated across broad bacterial diversity and *Pelagibacter* proteins are characteristically divergent, we applied a relaxed acceptance criterion of 0.75× the profile-specific threshold. The sensitivity of pathway results to this choice was assessed by repeating the analysis at 0.70× and 0.80×. As a negative control, the same relaxed pipeline was applied to the *E. coli* K-12 MG1655 proteome (GCF_000005845; NCBI-annotated proteins, 4,300 sequences). KEGG module completeness was computed as the fraction of module-defining KOs present per genome. Phylogenetic signal in variable pathway presence/absence was tested using a custom Fitch parsimony implementation in Python (tree parsed with BioPython v1.84): for each binary trait (pathway step present/absent), the minimum number of state changes on the unrooted 135-genome phylogeny was computed and compared to a null distribution generated by randomly permuting trait values across tips (10,000 permutations, random seed 42). P-values were computed as the fraction of permutations with equal or fewer changes than observed and Bonferroni-corrected for 5 tests.

### Structural annotation

Protein structures were predicted for 26,941 proteins (24,025 cluster representatives plus additional proteins from an initial ESMFold run) using ESMFold v2.0 (Lin et al., 2023) on an NVIDIA RTX 6000 Ada Generation GPU. Predicted structures were searched against the AlphaFold Database v4 (Varadi et al., 2024) using Foldseek v10.941 (van Kempen et al., 2024) with e-value ≤ 10⁻³, sensitivity 9.5, structural alignment mode (`--alignment-type 2`), and up to 5 hits per query. The query database was created with `foldseek createdb` and searched with `foldseek search` using 128 threads.

### Singleton cluster characterization

Singleton clusters (present in exactly one genome) were characterized by cross-referencing cluster representative proteins against the UniRef90 annotations. Functional categories were assigned by keyword matching on UniRef90 descriptions. Keywords: phage (phage, capsid, tail, terminase, portal, baseplate, head), mobile elements (transposase, integrase, recombinase, insertion), restriction-modification (restriction, methyltransferase, methylase, hsdM/R/S), toxin-antitoxin (toxin, antitoxin, vapB/C, mazE/F, relB/E), transport (transporter, permease, ABC, TRAP, efflux, pump), cell surface (glycosyltransferase, glycoside hydrolase, polysaccharide, capsular, lipopolysaccharide, O-antigen, wzy, wzx), and hypothetical (hypothetical, uncharacterized, DUF, domain of unknown function). Proteins matching none of these keywords but having a UniRef90 hit were classified as “other annotated”; those without any hit were classified as “no hit”. Per-gene GC content was calculated from the nucleotide sequences (.ffn files). Genomic islands were detected as maximal runs of ≥2 consecutive singleton genes in positional order along each chromosome (any strand), using the GFF coordinate annotations. A run was broken by any intervening non-singleton gene; no maximum intergenic distance was imposed since *Pelagibacter* intergenic spaces are uniformly short (median 4 bp). To trace the taxonomic origin of horizontally acquired singletons, the Tax= field was parsed from UniRef90 hit descriptions and classified by phylogenetic distance from *Pelagibacter*: same genus (Pelagibacter/SAR11), same order (Pelagibacterales), same class (Alphaproteobacteria), same phylum (other Proteobacteria), or cross-phylum. Candidate recent HGT events were identified as singletons with non-*Pelagibacter* best hits at ≥70% amino acid identity. Island mosaicism was assessed by examining the taxonomic origin of consecutive genes within individual genomic islands. The GC difference between singleton and non-singleton clusters was tested with a two-sided Mann-Whitney U test (SciPy v1.12.0).

### Hypervariable region and phage analysis

The hypervariable region (HVR) was defined by the chromosomal position of the largest singleton islands: because all genomes were reoriented to start at dnaA, the HVR was identified as the 7–15% window of each genome (measured from position 0). The presence and size of singleton islands in this window were tallied across all 135 genomes. To test whether HVR genes are carried by co-occurring phage, HVR singleton protein sequences from each sample’s *Pelagibacter* genomes were searched against viral protein sequences from the same sample (identified by geNomad v1.11.2 `end-to-end`, database v1.9, default score thresholds; Camargo et al., 2024) using MMseqs2 `easy-search` (e-value ≤ 10⁻³, top 5 hits, 2 threads per sample). Matches to geNomad-identified proviruses (integrated prophages) were tallied separately. HVR boundary conservation was assessed by examining the identity and position of tRNA genes (from tRNAscan-SE) and rRNA genes in the flanking regions (5–7% and 15–17% from dnaA). rRNA genes were identified using Infernal v1.1.5 (Nawrocki & Eddy, 2013) cmsearch against three Rfam covariance models: RF00177 (bacterial SSU/16S), RF02541 (bacterial LSU/23S), and RF00001 (5S). All 135 rotated genomes were searched with 32 CPUs and –noali. Chromosomal positions were expressed as fractions of genome length from dnaA, and the ITS spacer between 16S and 23S and the distance to 5S were computed from hit coordinates to assess operon structure. To test for an internal age gradient, each HVR singleton gene’s position was normalized to 0 (origin-proximal boundary) to 1 (terminus-proximal boundary) within the 7–15% window. GC content, deviation from the genome mean GC, and UniRef90 sequence identity were plotted against normalized position and tested with Spearman rank correlation across all HVR singletons pooled from 132 genomes. Smoothed profiles were computed by binning genes into 50 equal-width position bins and computing the mean per bin. As a negative control for phage-HVR matching, non-HVR non-singleton proteins from the same sample were searched against the same viral protein database, and match rates were compared across identity thresholds.

### Genome synteny

Gene order was compared across all 135 genomes using the protein cluster assignments from the pangenome analysis. For each genome, the ordered sequence of cluster IDs (with strand) was extracted from the GFF annotation. Gene adjacencies were defined as consecutive pairs of clustered genes in positional order along the chromosome; unclustered proteins were excluded. The adjacency frequency spectrum was computed by counting how many genomes shared each unique adjacency. Pairwise synteny similarity between genomes was quantified as the Jaccard index (shared adjacencies / union of adjacencies). The relationship between synteny conservation and sequence divergence was assessed by plotting pairwise Jaccard similarity against pairwise ANI for 500 randomly sampled genome pairs.

Operons were predicted for each genome using two criteria applied to consecutive genes: (1) co-directionality (same strand) and (2) intergenic distance <100 bp. Runs of two or more genes meeting both criteria were classified as predicted operons. Operon adjacencies (consecutive gene pairs within a predicted operon) were tallied across all 135 genomes to identify conserved co-transcriptional units. Conserved pairwise adjacencies (present in ≥120 genomes) were chained into maximal blocks by following unambiguous directed edges: starting from a node with no conserved predecessor (or multiple predecessors), the chain extends through nodes that have exactly one conserved successor whose sole predecessor is the current node. Proteins from conserved adjacencies were annotated by MMseqs2 `easy-search` against UniRef90 (UniProt Consortium, 2024) with an e-value threshold of 1 × 10⁻¹⁰, retaining the top hit per query.

### Transporter inventory

Transport-related KOs were extracted from the KofamScan results (relaxed threshold) and grouped by transport system (TRAP, ABC amino acid, ABC branched-chain amino acid, ABC iron(III), ABC glycine betaine/proline, etc.). Copy numbers were tallied per genome as the number of unique proteins assigned to each KO. To test for compensatory transporter expansion in auxotrophic genomes, substrate-specific transporter SBP copy numbers were compared between genomes with and without each variable biosynthetic pathway step using two-sided Mann-Whitney U tests, with Bonferroni correction for 3 tests (isoleucine/branched-chain AA SBP, histidine/polar AA SBP, serine/polar AA SBP). Pantothenate and NAD lack specific characterized transporters and were not tested.

### Sequencing depth comparison

The 0S assembly (station 8, summer, 6 flow cells total: the original 2 plus 4 additional) was compared to the 8S assembly (same station and date, 2 flow cells only) and all other standard-depth assemblies. Species found only in 0S were searched against all standard-depth assemblies using skani to assess fragment presence and coverage.

### NCBI genome comparison

All genomes classified as *Pelagibacter* (taxon ID 1655637) were downloaded from NCBI using the NCBI Datasets CLI v18.10.2 (n = 1,473; 1,442 after excluding the 31 already in the dataset). These were searched against a skani database built from our 135 genomes using `skani search` with the `--slow` preset and 2 threads. Best-hit ANI was used to classify each NCBI genome as same species (≥95% ANI), same genus (80–95%), or below detection (<80%). Assembly levels were extracted from the NCBI assembly report.

## Data availability

All genome sequences and annotations have been deposited at [repository TBD]. Assembly and annotation scripts are available at [repository TBD].

## Acknowledgments

We thank Erica Nejad and the crew of USGS R/V David H. Peterson for collecting samples. We cannot overemphasize the value of the support we have received from them. We also thank Miten Jain for lively discussions about nanopore sequencing and providing support for the PromethION sequencing.

Torben also wants to thank Lauren for putting up with how he does things; it can be challenging.

## Supplementary Information

**Supplementary Table 1:** Genome statistics for all 135 *Pelagibacter* genomes — Genome ID, Source, Genome size (bp), GC (%), Gene count, Coding density (%), Completeness (%), Contamination (%), GTDB-Tk classification, Species assignment.

**Supplementary Table 2:** Core gene list — 84 strictly core clusters with UniRef90 annotations, copy status (single-copy/paralog/truncated), and phylogeny inclusion.

**Supplementary Table 3:** KofamScan relaxed threshold validation — cross-reference of KO assignments at 0.75× threshold against independent UniRef90 annotations.

**Supplementary Table 4:** Variable pathway presence/absence by species cluster — 52 clusters × 5 variable pathways, showing fraction of genomes per species with each pathway step present.

**Supplementary Table 5:** Conserved operon blocks — 58 multi-gene units conserved across ≥120 of 135 genomes, with functional annotations.

**Supplementary Table 6:** NCBI accession numbers for the 31 *Pelagibacter* genomes downloaded from public databases.

**Figure S1:**
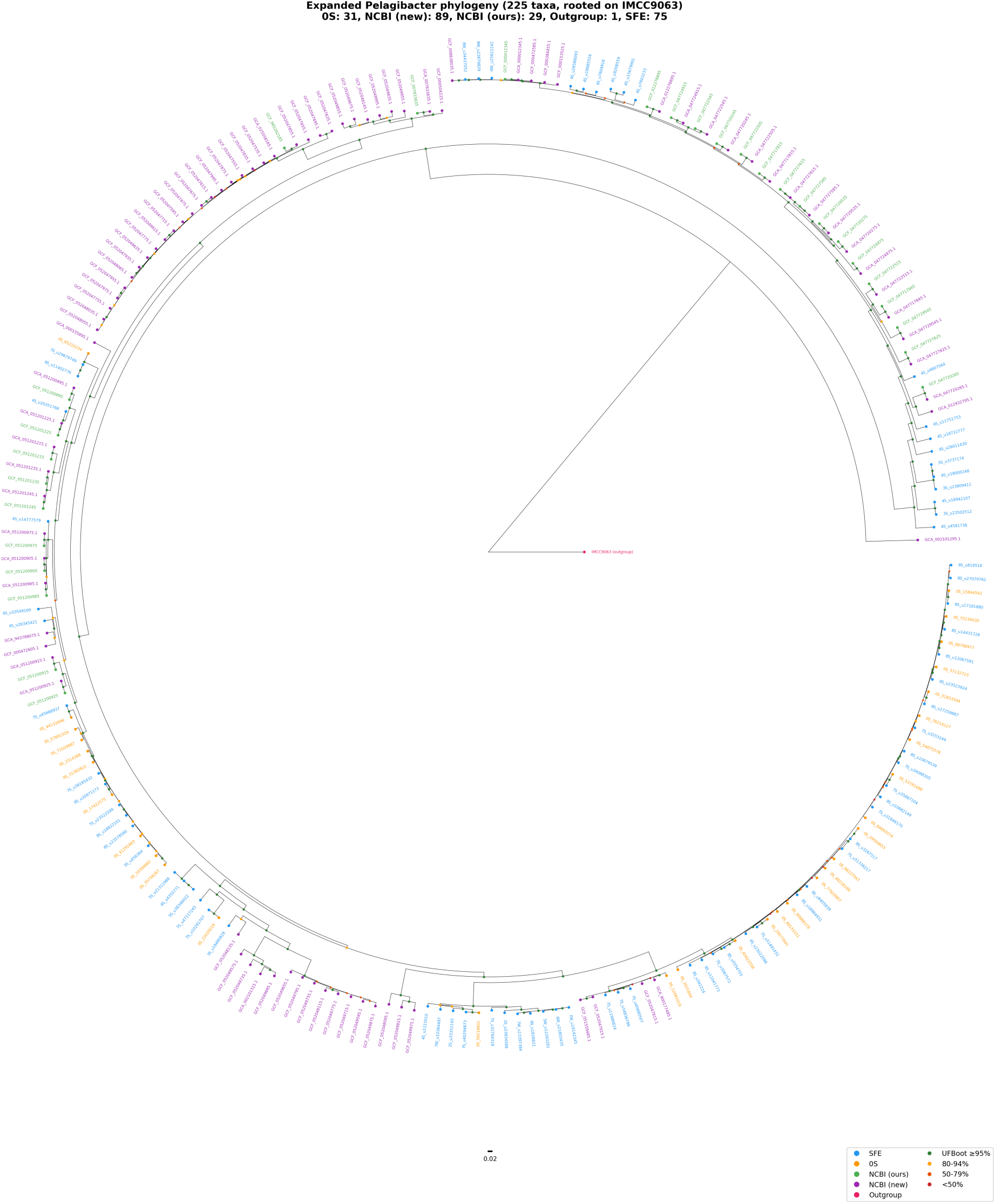
Expanded phylogenomic tree of 225 *Pelagibacter* genomes (135 from this study + 89 additional high-quality NCBI genomes + IMCC9063 outgroup). Tips colored by source. NCBI genomes nest within clades defined by SFE genomes rather than forming separate branches.

**Figure S2:**
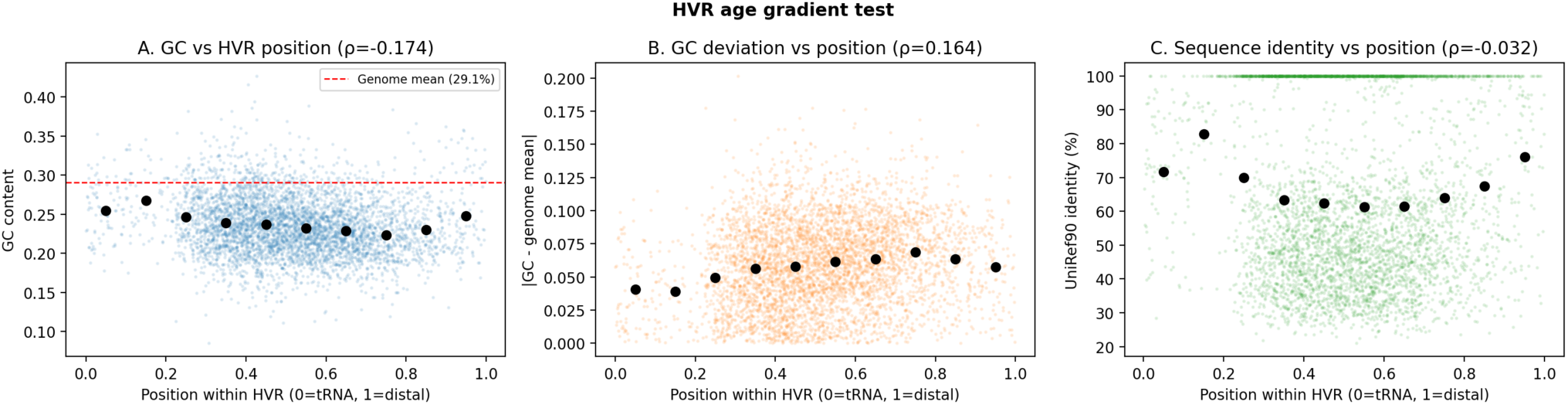
HVR internal structure. (A) GC vs position within HVR. (B) GC deviation from genome mean vs position. (C) UniRef90 identity vs position — U-shaped, highest at boundaries.

